# The loci of behavioral evolution: evidence that *Fas2* and *tilB* underlie differences in pupation site choice behavior between *Drosophila melanogaster* and *D. simulans*

**DOI:** 10.1101/494013

**Authors:** Alison Pischedda, Michael P. Shahandeh, Thomas L. Turner

## Abstract

The recent boom in genotype-phenotype studies has led to a greater understanding of the genetic architecture of a variety of traits. Among these traits, however, behaviors are still lacking, perhaps because they are complex and environmentally sensitive phenotypes, making them difficult to measure reliably for association studies. Here, we aim to fill this gap in knowledge with the results of a genetic screen for a complex behavioral difference, pupation site choice, between *Drosophila melanogaster* and *D. simulans*. In this study, we demonstrate a significant contribution of the X chromosome to the difference in pupation site choice behavior between these species. Using a panel of X-chromosome deletions, we screened the majority of the X chromosome for causal loci, and identified two regions that explain a large proportion of the X-effect. We then used gene disruptions and RNAi to demonstrate the substantial effects of a single gene within each region: *Fas2* and *tilB*. Finally, we show that differences in *tilB* expression correlate with the differences in pupation site choice behavior between species. Our results suggest that even complex, environmentally sensitive behaviors may evolve through changes to loci with large phenotypic effects.

## Introduction

The behaviors of closely related species can be remarkably different, and these differences can have important biological consequences. Behavioral evolution in insects has major impacts on crop decimation and disease vectoring (Karageorgi et al., 2017; McElhany et al., 1995). For example, one subspecies of the yellow fever mosquito, *Aedes aegypti*, prefers to bite humans, while another closely related species prefers other animals (McBride et al., 2014; Peterson, 1977). Understanding insect preferences therefore presents a major inroad to effective disease and pest management (Zhou et al., 2010). Behaviors are also critical to the creation and maintenance of biodiversity, as host, habitat, and mating preference behaviors are often key players in speciation and local adaptation (Coyne & Orr, 1997, 2004).

Despite the importance of behavioral traits, we know little about the genetic basis of their evolution. GePheBase, the most extensive compilation of natural genetic variants associated with trait differences, currently catalogs over 1800 associations, of which only 22 are for behavior (Martin & Orgogozo, 2013). From these 22, and others in the literature, it is clear that individual genes can sometimes have large effects on evolved differences in behavior (Leary et al., 2012; McGrath et al., 2011; Prince et al., 2017). It also clear that changes to sensory receptors in the peripheral nervous system can explain dramatic shifts in behavior (Cande et al., 2013; Leary et al., 2012; McBride et al., 2014). With so few studies, however, it is difficult to conclude how frequently we expect single loci to have large effects or how often sensory receptors explain species differences, due to pervasive ascertainment bias (Rockman, 2012). Indeed, a recent study discovered an exception to this emerging pattern: the evolution of a central neural circuit, rather than a peripheral sensory neuron, explains differences in mating behavior in two *Drosophila* species (Seeholzer et al., 2018).

The current lack of genetic studies mapping behavioral variation presumably arises from the fact that behaviors are difficult to measure reliably and repeatedly. Behavioral phenotypes often integrate multiple cues, are sometimes context dependent, and can be innate or learned, making it difficult to exclude environmentally induced variation. To better understand the genetic basis of behavioral evolution, we therefore need more case studies, with a focused effort on “metamodel” systems with documented behavioral differences between closely related species (Kopp, 2009).

Flies in the genus *Drosophila* are well poised to address these challenges. In *Drosophila*, hundreds of genetically identical individuals from variable wild-caught strains can be reared in a common environment, isolated at the beginning of their adult life stage, and repeatedly assayed for a trait of interest. Such a design significantly reduces the potential for environmentally induced variation to obscure genetic differences in behavior. Additionally, there are many *Drosophila* species with large, characterized differences in a variety of complex behaviors (Orgogozo & Stern, 2009). Undeniably, work comparing the repeated evolution of morphological traits across closely related species of *Drosophila* has significantly advanced our understanding of the general patterns linking genotype and phenotype for developmental traits (Kittelmann et al., 2017; Rebeiz & Williams, 2017; Sucena et al., 2003). Studies that investigate genetic complexity in *Drosophila*, where fine-mapping and functional follow-up are possible, should make similar progress for behaviors.

In the present study, we seek to address the lack of behavioral association studies using the *Drosophila* model system. Here, we investigate the difference in pupation site preference between two *Drosophila* species: *D. melanogaster* and *D. simulans*. When the larval stages of these species are ready to metamorphose into adults, they first enter a pupal stage. The pupa, which lasts for a number of days, is immobile and therefore vulnerable to parasitism, predation, desiccation, and disease (Markow, 1981a). Before pupating, larvae enter a “wandering” stage, where they search for an appropriate pupation site (Riedl et al., 2007; Sokolowski et al., 1984). Depending on the strain and species, larvae vary from pupating directly on their larval food source to traveling more than 40 cm away from it (Stamps et al., 2005). This behavior has been extensively studied, and is exquisitely sensitive to environmental conditions— individuals alter their behavior in response to light, moisture, pH, the presence of other species, parasitism, and more (Hodge & Caslaw, 1998; Markow, 1981b; Sameoto & Miller, 1968a; Seyahooei et al., 2009). Despite this environmentally induced variation, the effects of genotype on preference are considerable. Within species, strains and populations often differ in how far they travel from their food source before pupating, although the most consistent experimentally demonstrated differences are between species (Markow, 1979; Vandal et al., 2008). Interestingly, differences in pupation site choice behavior between species do not correspond to their taxonomic classification (Shivanna et al., 1996). For example, *D. melanogaster* and *D. simulans* shared a common ancestor 2-3 million years ago (Lachaise & Silvain, 2004), and are extremely similar in terms of their ecology, morphology, and physiology (Parsons, 1975). Previous work shows, however, that they differ markedly in terms of pupation site choice, with *D. simulans* pupating closer to the larval food source, on average (Markow, 1979, 1981a). This difference is not due to laboratory adaptation, as freshly collected individuals show the same pattern (Markow, 1979, 1981a). These species are frequently collected in the same microhabitats, and their differences in pupation site choice behavior have been postulated to be a form of niche partitioning (Markow, 1979, 1981a). Supporting this hypothesis, pupation site choice responds to density dependent selection in the laboratory (Mueller & Sweet, 1986), and provides a potential increase in competitive ability between species ovipositing in the same media (Arthur & Middlecote, 1984).

Here, we investigate the genetic basis of this difference in pupation site choice between *D. melanogaster* and *D. simulans*. Despite substantial reproductive isolation, *D. melanogaster* females can be persuaded to mate with *D. simulans* males in the lab, and vigorous female F1 hybrids result from the cross. Males are usually inviable, but we use a *D. simulans* hybrid male rescue strain (Brideau et al., 2006a; Watanabe, 1979) to circumvent this challenge, and show that a significant proportion of the species difference in pupation behavior can be mapped to the X chromosome, consistent with findings using other *Drosophila* species (Erezyilmaz & Stern, 2013). Still, these hybrids remain sterile, so genetic dissection using a backcross mapping population is not possible. Instead, we use a widely available set of transgenic *D. melanogaster* deficiency lines to screen a substantial portion of the X chromosome (Cook et al., 2012; Ryder et al., 2004). We use these lines to create hybrid females that lack large, overlapping portions of the X chromosome from *D. melanogaster*, and therefore express only *D. simulans* alleles in those regions. We identify two loci with large effects on pupation behavior. We then employ genetic knockouts of candidate loci within these regions to demonstrate their effects, and use RNAi knockdown to further test the role of two genes, *touch insensitive larvae B* (*tilB*) and *Fasciclin 2* (*Fas2*). Finally, we use real time qRT-PCR to test for species-level differences in gene expression of *tilB* in larvae, and show that *tilB* is more highly expressed in *D. melanogaster* than in *D. simulans*.

## Methods

### General fly maintenance

Unless otherwise stated, we maintained all fly strains for these experiments in 20 mm diameter vials containing standard cornmeal-molasses-yeast medium at 25°C under a 12h:12h light/dark cycle at 50% relative humidity. Under these conditions, we established non-overlapping two-week lifecycles as follows. For all stocks, except LH_M_ and *Lhr* (see below), we transferred male and female adult flies into fresh vials containing food media supplemented with live yeast on the surface for 1-3 days, at which point the flies were discarded. 14 days later (after all progeny had eclosed), we again transferred adult flies into fresh vials for 1-3 days to begin the next generation. We maintained LH_M_ and *Lhr* identically, except we additionally regulated density by transferring only 10 males and 10 females to begin the next generation.

### Characterizing pupation behavior for *D. melanogaster* and *D. simulans*

We measured pupation behavior for 11 *D. melanogaster* and 12 *D. simulans* strains collected from various locations throughout the world (Table S1). The 11 *D. melanogaster* strains included 10 of the “founder” wild-type inbred lines of the Drosophila Synthetic Population Resource (King et al., 2012; flyrils.org), and a single wild-type line created from the LH_M_ laboratory-adapted population (Rice et al., 2005). The 12 *D. simulans* strains included 11 wild-type strains and a single strain carrying *Lethal hybrid rescue* (*Lhr*), a mutation that restores viability in *D. melanogaster*/*D. simulans* hybrid males.

To measure pupation behavior, we placed 10 males and 10 females from a specific line (both 3-5 days old) into half pint bottles and allowed females to oviposit overnight on a 10 mm diameter petri dish filled with food medium that was placed in the opening of the bottle. In total, we set up 5 bottles for each line. The following morning, we transferred 100 eggs from the petri dishes into vials containing food medium that were lined with an acetate sleeve on which the larvae could pupate. In total, we set up 5-8 vials per line. Vials were held at 25°C for 8 days, at which time the liner was removed and the locations of the pupae were recorded (8 days was long enough for almost all larvae to pupate without any flies eclosing). A pupa was considered “on” the food if it was within 1 cm of the food surface, while all pupae that were further than 1 cm from the food surface were considered “off” the food. For each vial, we calculated the proportion of pupae on the food surface. For comparisons between species, our unit of replication was the mean proportion of pupae on the food surface from each line (i.e. N=11 for *D. melanogaster* and N=12 for *D. simulans*).

### Crossing D. melanogaster with D. simulans

For all crosses below, we created F1 hybrids between *D. melanogaster* and *D. simulans* using the following protocol. *D. simulans* males were collected as virgins within 6 hours of eclosion and held at room temperature in groups of 20 in vials containing food medium for 3-4 days. To set up crosses, we collected young *D. melanogaster* virgin females within 2-3 hours of eclosion, and combined 8-12 of these females with 20 *D. simulans* males in vials containing food medium supplemented with an *ad lib* amount of live yeast on the surface. We then pushed a long foam plug down into the vial, leaving approximately 1 cm of space above the food surface. We held flies under these conditions for 3 days, at which time they were transferred from these “cross vials” into “pupation vials” that contained food medium with no added yeast, and were lined with an acetate sleeve on which the larvae could pupate. We always set up crosses using *D. melanogaster* females and *D. simulans* males, because crosses in the opposite direction were never successful.

### Measuring pupation behavior in F1 hybrids

To create F1 hybrid males and females, we used *D. simulans* males from the *Lethal hybrid rescue* (*Lhr*) strain (Watanabe, 1979). The *Lhr* mutation restores viability in F1 hybrid males, which are usually lethal (Brideau et al., 2006b). To create F1 hybrid females and F1 males with a *D. melanogaster* X chromosome (“melX” males), we crossed wild-type females from our LH_M_ strain (provided by William Rice) to *Lhr D. simulans* males (Fig 1A). Because we were unable to successfully cross *D. simulans* females to *D. melanogaster* males, we created F1 hybrid males with the *D. simulans* X chromosome (“simX” males) by crossing *D. melanogaster* LH_M_ females that carry a compound X chromosome (*C(1)DX y f*) (Rice et al., 2005) to *D. simulans Lhr* males. The background of this strain is identical to that of LH_M_, with the exception of the compound X chromosome. The compound X in these females ensured that the X chromosome was transmitted from *D. simulans* fathers to their F1 hybrid sons (Fig 1B). For each direction of the cross, we combined *D. melanogaster* females with *D. simulans* males, as described above. This crossing scheme ensures that all maternal inheritance (cytoplasmic and mitochondrial) in the reciprocal male hybrid crosses originates from the *D. melanogaster* parent. Thus, these hybrids have an identical background with the exception of the sex chromosomes, and any differences we observe between melX and simX males are directly attributable to their different sex chromosomes.

**Fig 1.**
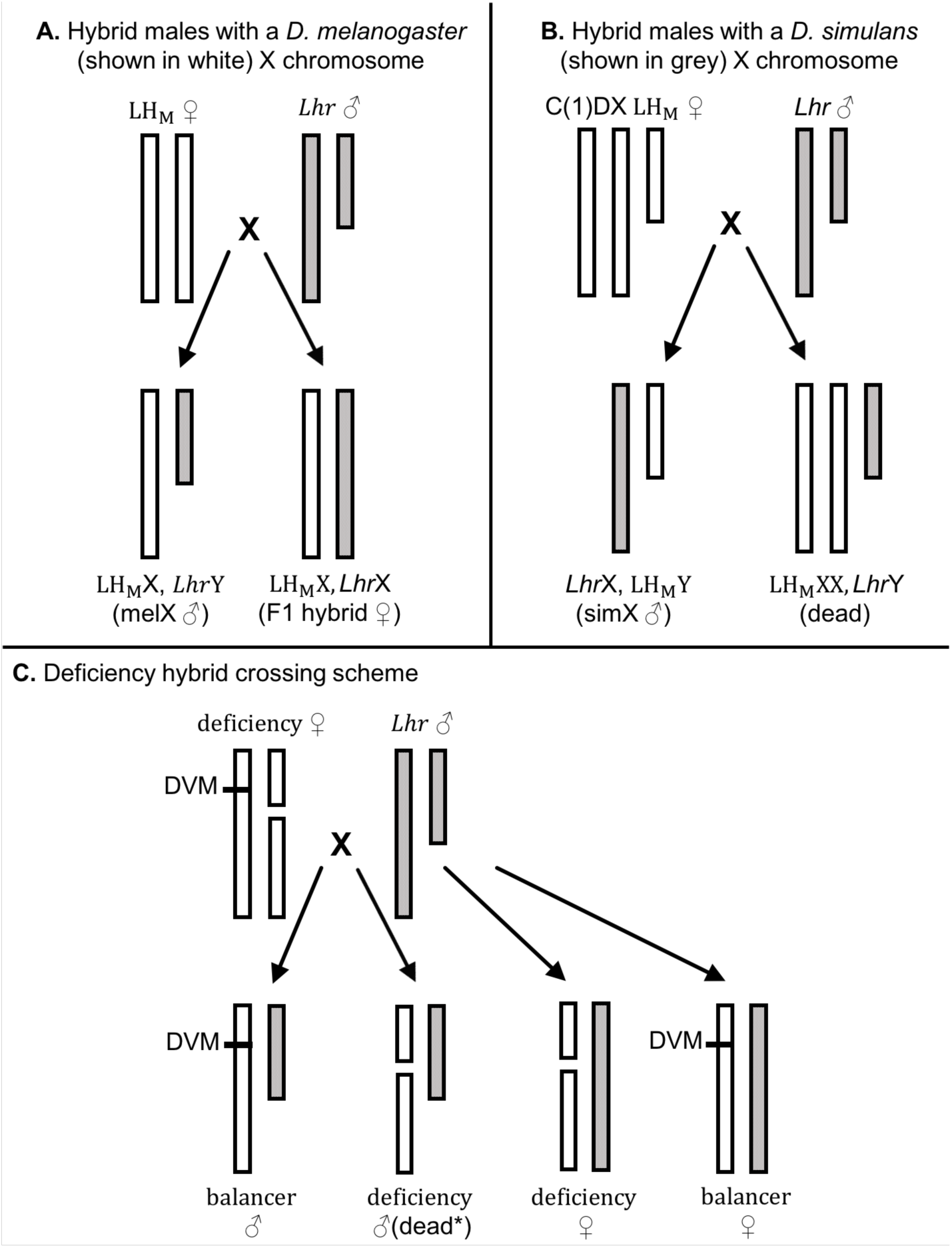
Crossing schemes to generate reciprocal X chromosome hybrid males and deficiency/balancer hybrid females. **A.** Crossing wild-type *D. melanogaster* females (LH_M_, shown in white) to *D. simulans* males (*Lhr*, shown in grey) produces hybrid males with a *D. melanogaster* X chromosome (melX) and hybrid females. **B.** Crossing a compound X C(1)DX LH_M_ female to *Lhr* males produces hybrid males with a *D. simulans* X chromosome (simX). Females of this cross would inherit two *D. melanogaster* X chromosomes and a *D. simulans* Y chromosome, but are inviable. **A and B.** Note that the background of the reciprocal male hybrids resulting from each cross (melX and simX) is an identical combination of *Lhr* and LH_M_ with the exception of the sex chromosomes. **C.** Crossing *D. melanogaster* X chromosome deficiency lines, which have a balancer X chromosome with a dominant visible marker (DVM) and an X chromosome with a large deletion, to *Lhr* produces deficiency hybrid females, balancer females, balancer males, and deficiency males (*mostly dead due to large deletions on a hemizygous chromosome with some deficiency lines being exceptions).

After 3 days in the cross vial, we transferred males and females into pupation vials for 24 hours, at which time the flies were removed. While screening hybrid pupation behavior, we also concurrently screened pupation behavior for the parental *D. melanogaster* strain (LH_M_) and the parental *D. simulans* strain (*Lhr*) for comparison. Parental strain cross vials contained only a moderate amount of yeast, were set up with only 5 males and 5 females (pure species crosses produce more offspring), and did not have a plug pushed down into the vial, but were otherwise treated identically to the hybrid crosses. In total, we set up 30-33 vials per treatment.

All pupation vials were held at 25°C for 8 days, at which time the liner was removed. We removed any remaining larvae, and cut the liner at a point 1 cm above the food surface. The portion of the liner that contained pupae within 1 cm of the food surface was returned to the original vial (the “on vial”), while the portion of the liner with pupae further off of the food surface was placed in another vial containing food medium (the “off vial”). The flies that eclosed were sexed and counted 7 days later (15 days post-egg); all flies that eclosed in the “on vial” were considered flies that pupated on the food surface, while all flies that eclosed in the “off vial” were considered flies that pupated off the food surface. We then calculated the proportion of individuals that pupated on the food for each type of individual (genotype and sex).

To assess the validity of using hybrid behavior to map interspecific differences, we set up an additional experiment to ensure that hybrids behaved typically with respect to pupation site choice behavior. This is a potential concern, as hybrids between *D. melanogaster* and *D. simulans* are known to differ from either parent in a variety of traits (Barbash & Ashburner, 2003; Sturtevant, 1920; Takano, 1998). For a subset of pupation vials, we calculated the average pupation height of males and females from each strain as follows. Instead of dividing the pupation liner into “on” food and “off” food sections, we cut the liner at 8 intervals, each spaced 1 cm from the last, starting at the food surface. These portions of the liner were ranked from 1 (on the food surface) to 8 (the furthest from the food surface), and were transferred to separate vials containing food medium. 7 days later the flies that eclosed were counted, and we used the rankings to calculate a mean pupation height, such that a higher value indicates a farther distance from the food. We performed this experiment for our *D. simulans* strain (*Lhr*), our reciprocal hybrid crosses (melX and simX in an LH_M_ background), the *D. melanogaster* strain w^1118^, and melX hybrids in a w^1118^ background (w^1118^ is the background strain for the majority of deficiencies we used in our screen).

### Mapping hybrid pupation behavior using the Bloomington Deficiency Kit

Because we found a significant effect of the X chromosome on the difference in pupation site choice behavior between *D. simulans* and *D. melanogaster*, we devised a crossing scheme using molecularly engineered chromosomal deficiencies to screen the X chromosome for loci contributing to this difference. These deficiencies are part of the Bloomington Deficiency Kit (Cook et al., 2012), available from the Bloomington Drosophila Stock Center (BDSC). We assayed a total of 90 deficiency strains covering 87% of the X chromosome (Table S2). We restricted our deficiency screen to lines from the BSC, Exelixis, and DrosDel sets to control for strain background effects while also maximizing chromosome coverage.

To set up crosses, we collected deficiency females as young virgins (2-3 hours after eclosing) and crossed them to *D. simulans* males from the *Lhr* strain. After 3 days in the cross vial, we transferred males and females into pupation vials for 24-48 h, at which time the flies were removed. We then divided the pupations vials into “on” and “off” vials as we did for our F1 hybrids (above).

These crosses produced two types of hybrid female that were heterozygous for *D. melanogaster*/*D. simulans* at each autosome (Fig 1C). The deficiency hybrid females contain the *D. melanogaster* deficiency X chromosome and a *D. simulans* X chromosome, making them hemizygous for a segment of the X chromosome. At this locus, these hybrid females only express *D. simulans* alleles. The balancer hybrid females contained the *D. melanogaster* balancer X chromosome (marked with a dominant visible marker) and a *D. simulans* X chromosome. These females are heterozygous for *D. melanogaster*/*D. simulans* over the entirety of the X chromosome, and thus express both *D. simulans* and *D. melanogaster* alleles. Although the deficiency hybrids are our flies of interest, the balancer hybrids provide an experimental control, as these females developed in the same environment as our experimental flies. As a result, we calculated the proportion of deficiency hybrid females that pupated on the food and the proportion of balancer hybrid females that pupated on the food. We then used these measures to calculate a “pupation index” as the proportion of deficiency hybrids pupating on the food divided by the proportion of balancer females pupating on the food. To increase the accuracy of our estimates, we only included pupation vials in our analysis that yielded at least 10 of each type of female. For each deficiency hybrid strain we measured, we report the median pupation index of all replicates, because there were often high-scoring outliers that significantly skewed the mean pupation index. These outliers almost always had abnormally high pupation indices, so focusing on median values makes our findings more conservative.

Any deficiency hybrid cross with a median pupation index greater than 1 indicates that more deficiency females pupated on the food compared to balancer females, potentially because the deficiency includes *D. melanogaster* genetic variation that is involved in pupation site choice behavior. Alternatively, simply creating flies that are hemizygous at a locus on the X chromosome may result in a variety of pleiotropic effects that make larvae less likely to climb up the vial. To test for this, when a deficiency hybrid cross showed a pupation index significantly greater than 1 (Table S2), we crossed that *D. melanogaster* deficiency strain to a *D. melanogaster* wild-type strain (T.4). If these *D. melanogaster* deficiency crosses displayed the same pattern, we considered the effect of the deficiency on pupation behavior to be a byproduct of deleting a large portion of the X chromosome, rather than revealing recessive *D. simulans* variation, and discarded them. If instead the pupation index for the *D. melanogaster* cross was significantly lower than the pupation index for the *D. simulans* cross, we pursued that deficiency for further validation. To ensure that this pattern is not a result of epistasis from the hemizygous region in a hybrid background, we further crossed these deficiencies to an additional *D. melanogaster* (T.1) and *D. simulans* (Mex180) strain to test for background-specific effects.

### Testing candidate genes in deficiency regions using gene knockouts and RNAi knockdown

For regions of interest identified by our deficiency screen, we ordered transgenic knockouts for any genes available within the region at the time. Our deficiency screen identified two regions of interest. The first is the overlap of Df(1)BSC869 and DF(1)ED6720, excluding the region covered by DF(1)ED6727, which did not have a pupation index greater than 1 (Fig 4A, Table S2). The resulting region of interest spans X:4,204,351 - 4,325,174 (Fig 4C). Within the first region, there are 23 genes, of which 20 are protein coding (Table S3A). According to modENCODE expression data, 15 of those 20 protein coding genes are expressed in larvae, while only 6 are also expressed in the larval nervous system, which we would expect for genes regulating behavior (Graveley et al., 2011). Five of these six genes are well described. At the time of assay, only three of the five characterized genes within this region had non-lethal verified loss-of-function alleles available: *Fas2* (Fasciclin2), *mei9* (meiotic 9), and *norpA* (no receptor potential A). We tested knockouts of each for an effect on pupation site choice behavior. We screened two *Fas2* knockouts, the mutant allele *Fas2^eb112^* (*Fas2^eb112^*/FM7c; Grenningloh et al., 1991; provided by Brian McCabe), and a p-element insertion allele, *Fas2^G0293^* (former BDSC Stock 11850; full genotype: *w^67c23^P{lacW}fas2^G0293^*/FM7c). It is important to note that the nature of the lesion is uncharacterized for *Fas2^G0293^* (although our behavioral data below suggest that it is indeed a loss of function allele, as it behaves indistinguishably from the verified knockout *Fas2^eb112^*). We additionally screened the *mei-9^A1^* mutant allele (*w^1^ mei-9^A1^*/FM7h; BDSC stock #6792) and the *norpA^36^* mutant allele (*w* norpA^36^*; BDSC stock #6792). Because *norpA^36^* is not held over a balancer, we also set up crosses using a *norpA* rescue strain created in the same background (*w* norpA^36^*; *P{w*[*+mC*]*=ninaE.norpA.E}2*; BDSC stock # 52276) as a control. It is worth noting that while *norpA* is expressed in the larval nervous system, it is only partially contained within this region, and is also largely deleted by another deficiency (Df(1)ED6727; Table S2) that did not display a pupation index significantly greater than 1 or 0.88 (the grand mean for all deficiency hybrids).

The second region of interest identified by our deficiency screen was the region deleted by DF(1)Exel6255 (X:21,519,203 – 22,517,665; Fig 4D). Within this region are 28 genes, of which 22 are protein coding (Table S3B). Of the 22 protein coding genes, 14 are expressed in larvae – 13 of which have some expression in the larval nervous system (Graveley et al., 2011). Of these, 7 are described. We obtained knockout strains for both of the characterized genes expressed in the larval nervous system that had verified loss-of-function alleles available at the time: *tilB* (touch insensitive larva B) and *wap* (wings apart). We screened two *tilB* mutant alleles, *tilB^1^* and *tilB^2^* (*y w tilB^1/2^* /FM4; Kernan et al., 1994; provided by Daniel Eberl), and the *wap^2^* mutant allele (*wap^2^*/FM6; BDSC stock # 8133).

Like the deficiency strains, each of our gene disruptions (with the exception of *norpA*) is held over a balancer chromosome with a visible marker. To measure the pupation behavior of hybrids containing knockout copies of these *D. melanogaster* genes, we crossed each *D. melanogaster* knockout strain to *Lhr* using the previously described methods, and calculated the pupation index as the proportion of knockout females on food / the proportion of balancer females on the food. As for our deficiency screen, any hybrid knockouts with a median PI greater than 1 for the hybrid cross suggest that the knockout gene may be involved in pupation site choice. We also crossed each *D. melanogaster* knockout strain to a wild-type *D. melanogaster* (T.4) strain to ensure that hybrid knockouts with a pupation index greater than 1 are not simply an artifact of being hemizygous for this particular gene. We crossed knockout strains that displayed the pattern we expect for a gene involved in pupation site choice (i.e. a pupation index significantly greater than 1 when crossed to *D. simulans*, which is also significantly greater than the pupation index when crossed to *D. melanogaster*) to an additional *D. simulans* (Mex180) and *D. melanogaster* (T.1) wild-type strain for verification. For *norpA*, we crossed the mutant and control strains to *Lhr*, and compared hybrid *norpA* mutant females to hybrid *norpA* rescue females. We additionally crossed these strains to the *D. melanogaster* T.4 strain.

Our knockout screen identified two genes that appear to be involved in pupation site choice: *tilB* and *Fas2*. We further tested the effects of these genes on pupation behavior using RNAi knockdown in *D. melanogaster*. We used the elav-Gal4 driver (P{w[+mC]=GAL4-elav.L}2/CyO; BDSC #8765), which expresses Gal4 throughout the nervous system. We drove down the expression of *tilB* and *Fas2* throughout the nervous system by crossing elav-Gal4 virgin females to UAS-*tilB* (BDSC #29391: y[1] v[1]; P{y[+t7.7] v[+t1.8]=TRiP.JF03324}attP2) and UAS-*Fas2* (BDSC #34084: y[1] sc[*] v[1]; P{y[+t7.7] v[+t1.8]=TRiP.HMS01098}attP2) males, respectively. The resulting flies express a gene-specific hairpin RNA throughout the nervous system, causing the degradation of mRNA, and thus, reduced expression of that gene (Perkins et al., 2015; Zeng et al., 2015). As experimental controls for each RNAi cross, we also crossed elav-Gal4 virgin females to the RNAi background stock (*y v; attP2, y+* (y^1^ v^1^; P{y[+t7.7]=CaryP}attP2; BDSC stock # 36303) and the Gal4-1 stock (containing a hairpin RNA targeting Gal4 in VALIUM20; BDSC stock # 35784). Together, these controls allow us to account for the effect of both the Gal4 mutation and general expression of hairpin RNA throughout the nervous system. Any differences we detect between these controls and our RNAi crosses must therefore be due to the expression of the gene-specific (*tilB* or *Fas2*) hairpin RNA. We set up pupation vials using the methods described above, and for each cross, we calculated the proportion of RNAi (or control) flies on the food (removing any data points with fewer than 20 experimental flies). If more RNAi flies pupate on the food in the experimental cross (in which the expression of the gene is driven down) compared to the control crosses (in which gene expression is unaffected), this provides further support for that gene’s involvement in pupation site choice. For these vials, we additionally determined the average pupation height of flies from each cross using the methods described above for F1 hybrids; if RNAi flies pupate closer to the food than flies from the control crosses, this suggests that gene may be involved in pupation site choice.

### Testing *tilB* for species-specific differences in larval transcript expression

We selected two each of our 11 *D. melanogaster* and 12 *D. simulans* strains to test for larval stage-specific expression differences of candidate genes using real time qRT-PCR – one extreme and one average. For *D. simulans*, we selected Geo288 and Per005 (Table S1), because Geo288 has the highest proportion of pupae on the food of the *D. simulans* strains and Per005 is closest to the species mean (Fig 2). For *D. melanogaster*, we selected CA1 and T.4 (Table S1) because T.4 has the lowest proportion of pupae on the food of the *D. melanogaster* strains, and CA1 is closer to the species mean (Fig 2).

**Fig 2.**
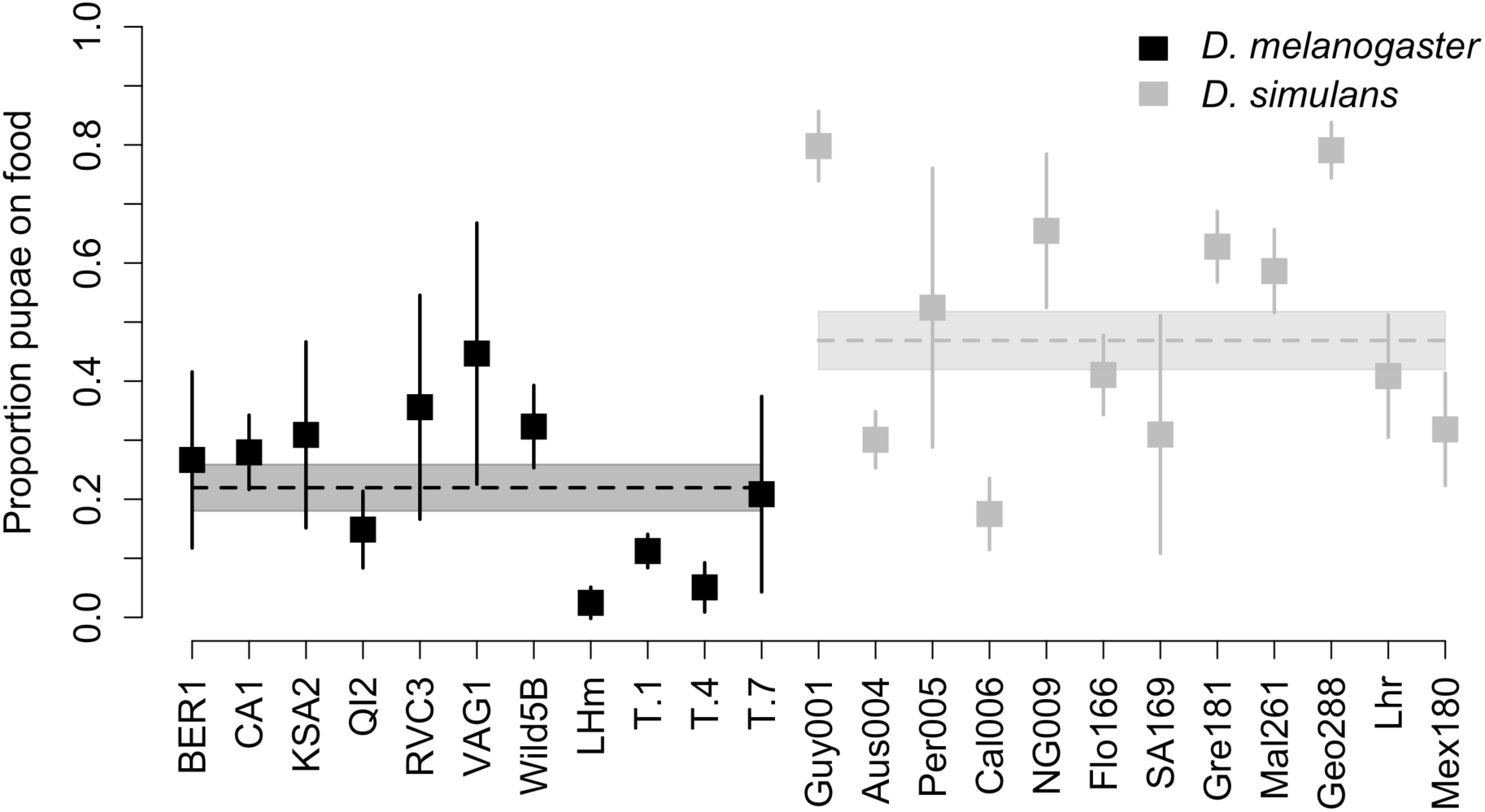
Pupation behavior differences between *D. melanogaster* and *D. simulans*. The mean proportion of individuals in a vial that pupated on the surface of the food for 11 *D. melanogaster* lines and 12 *D. simulans* lines described in Table S1. Error bars denote the 95% confidence interval around each individual mean (N = 5-8). The dashed horizontal lines indicate the grand mean for each species. The boxes surrounding the dashed lines denote the 95% confidence interval around the grand mean.

To harvest larvae from these strains, we allowed adult females to oviposit in standard vials containing food media between the hours of 8 AM and 12 PM over two consecutive days. 120 hours after the final oviposition day, we floated larvae out of the food media using a 20% sucrose in water solution, sucked them up using a transfer pipet, briefly rinsed them with DI water on cheesecloth, and snap froze them using liquid nitrogen (Nichols et al., 2012). In this way, we collected 20-30 mg of larvae from two developmental time points: 96 and 120 hours following oviposition. These time points approximate early wandering and late wandering larval stages. We chose these time points because they are presumably when pupation site choice occurs, and because the larvae are large enough for many to be harvested at once using the above methods. We extracted mRNA using the Qiagen RNeasy Plus Mini Kit, and prepared cDNA using the Promega Verso kit.

To quantify transcript abundance, we designed primers that span a single intron near the 3’ end of *tilB* (Huggett et al., 2005). Additionally, we used primers for the gene *RpL32* as an internal control (Al-Atia et al., 1985; Chertemps et al., 2007). *Fas2* is a complex gene with multiple splice forms, so we were unsuccessful in designing general primers that would amplify all transcripts in both species. For this reason, we did not include *Fas2* in these experiments. A full list of primers and transcripts can be found in Table S4A. For each stage and strain, we prepared two to three biological replicates, which we then amplified in two technical replicates for 40 rounds of qPCR. Using *RpL32* transcript number as an internal control, we calculated relative transcript abundance while correcting for species differences in primer efficiency. We estimated primer efficiency differences by serially diluting gDNA from each of our *D. melanogaster* and *D. simulans* strains, performing qPCR, and using a standard curve to calculate adjusted amplification factors (Table S4B). To ensure that we were amplifying cDNA made from RNA, and not gDNA contamination, we performed gel electrophoresis on our cDNA samples to ensure we only visualized the short, intron-less, band.

### Estimating effect sizes

We used our results to estimate how much of the difference in pupation site preference between *D. melanogaster* and *D. simulans* can be attributed to: *i*) the X chromosome, *ii*) our deficiencies of interest (Df(1)BSC869/DF(1)ED6720 and Df(1)Exel6255), and *iii*) *Fas2* and *tilB*. We first calculated the “species difference ratio” (using the data from our parental/hybrid screen in Fig 3) by dividing the median proportion of males on the food for *D. simulans* by the median proportion of males on the food for *D. melanogaster* males (species difference ratio = 5.27). We then calculated an “X effect ratio” by dividing the median proportion of simX males on the food by the median proportion of melX males on the food. To determine how much of this species difference can be attributed to the X chromosome, we divided the “X effect ratio” by the “species difference ratio”.

**Fig 3.**
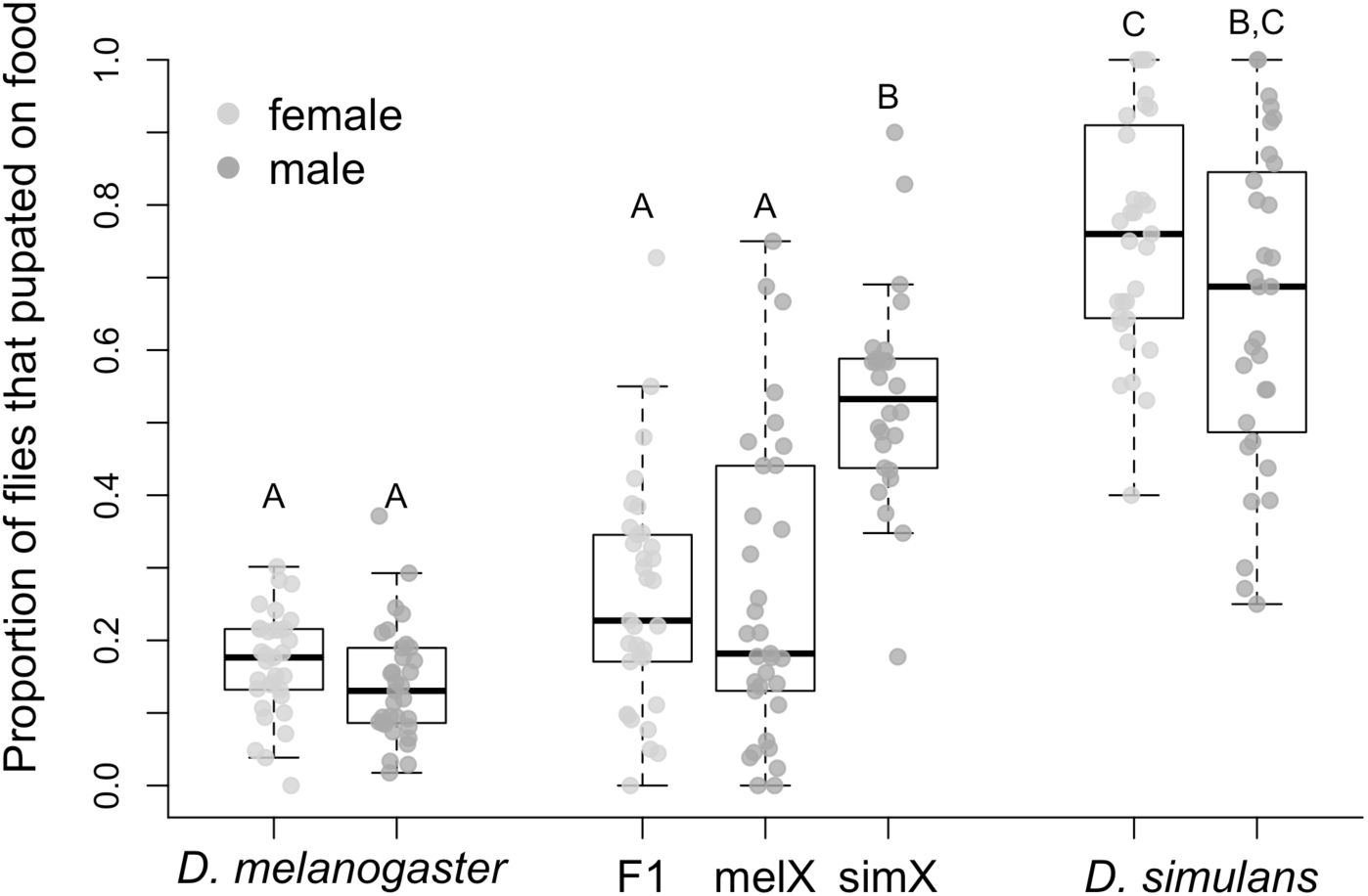
Pupation site choice behavior for *D. melanogaster*, *D. simulans*, and their F1 hybrids. The proportion of individuals that pupated on the food surface for males and females from both species and their F1 hybrids. *D. melanogaster* males and females were taken from the LH_M_ strain, while *D. simulans* males and females were taken from the *Lhr* strain. F1 hybrids resulted from a cross between these two strains. The “melX” hybrid males have the *D. melanogaster* X chromosome, and the “simX” hybrid males have the *D. simulans* X chromosome. Both hybrids have *D. melanogaster* cytoplasmic inheritance. Box plots display the median (bold bar), interquartile range (box), and full extent of the data set excluding outliers (whiskers). Bars labeled with different letters are significantly different from one another after sequential Bonferroni correction for multiple comparisons (p < 0.0001 after correction, N = 26-33).

We then estimated the effect size of our deficiencies by calculating “deficiency effect ratios” (the median pupation index when the deficiency was crossed to the *D. simulans Lhr* strain/the median pupation index when the deficiency was crossed to the *D. melanogaster* T.4 strain). Because Df(1)BSC869 and DF(1)ED6720 overlap, we used their mean deficiency effect ratio to estimate the effect size of the overlapping region. To determine how much of the X effect difference can be attributed to each deficiency region, we divided the “deficiency effect ratio” by the “X effect ratio”.

Finally, we estimated the effect size of our two identified candidate genes, *Fas2* and *tilB*. For each allele we tested, we calculated a “knockout effect ratio” (the median pupation index when the knockout was crossed to the *D. simulans Lhr* strain/the median pupation index when the knockout was crossed to the *D. melanogaster* T.4 strain), and then we used the average knockout effect ratio of the two alleles for each gene (i.e. *Fas2*: average of *Fas2^eb112^* and *Fas2^G0293^*; *tilB*: average of *tilB^1^* and *tilB^2^*). We then determined the contribution of each gene to its deficiency effect size, and to the overall X effect, by dividing the “knockout effect ratio” by the “deficiency effect ratio” and the “X effect ratio”, respectively.

For all effect size estimates, we calculated bootstrapped 95% confidence intervals using 100,000 bootstraps.

### Data availability

All relevant data supporting the findings of this study are represented in the figures and supplementary tables.

## Results

### 1. Differences in pupation behavior between *D. melanogaster* and *D. simulans*

We found significant variation among the 11 *D. melanogaster* and 12 *D. simulans* strains in the proportion of individuals that pupated on the food surface (Fig 2; *D. melanogaster* Wilcoxon test: χ^2^= 42.69, df = 10, p<0.0001; *D. simulans* Wilcoxon test: χ^2^= 56.34, df = 11, p<0.0001). When we tested for a species difference in pupation behavior, we found that our *D. simulans* lines had significantly more pupae on the food surface compared to our *D. melanogaster* lines, on average (Fig 2; Wilcoxon test: χ^2^= 8.37, df = 1, p = 0.0038).

Although we controlled egg density to characterize species differences in pupation behavior, there were viability differences among our surveyed lines. As a result, we had significant variation in the total number of pupae in each vial for our *D. melanogaster* lines (ANOVA: F_10,65_=5.58, p<0.0001) and our *D. simulans* lines (ANOVA: F_11,66_=8.01, p<0.0001). Previous studies have found that larval density correlates with pupation height in *D. melanogaster* (Sokal et al., 1960). To control for differences in density, we also performed the same analyses above using the residuals from a regression between number of pupae in the vial and proportion of pupae on the food. None of our findings changed using this analysis (Fig S1), indicating that our results were not affected by variation in larval density.

Although most of our *D. melanogaster* lines have been in the lab since the 1950s and 1960s, we found no difference in the proportion of pupae on the food between *D. simulans* strains collected in the 1950s/1960s and those collected in the 2000s (Table S1; Wilcoxon test: χ^2^= 0.24, df=1, p=0.62), indicating that the species difference we report here is unlikely to be an artifact of laboratory adaptation in our surveyed strains.

### 2. Differences in pupation behavior have a significant X effect

We found significant differences among F1 hybrid genotypes when we screened hybrids alongside their parental strains (Fig 3; Full model Wilcoxon test: χ^2^= 144.57, df= 6, p<0.0001). Specifically, a significantly higher proportion of F1 hybrid males pupated on the food when they had inherited a *D. simulans* X chromosome (simX males) compared to a *D. melanogaster* X chromosome (melX males), indicating that this species divergence in pupation behavior has a significant X effect (p<0.0001 after correcting for multiple comparisons; Fig 3). This is supported by the fact that the proportion of individuals that pupated on the food was not significantly different between melX hybrid males and *D. melanogaster* males, or between simX hybrid males and *D. simulans* males. All of these findings are unchanged when we control for density effects (Fig S2).

Additionally, we found no difference in the proportion of F1 hybrid females and melX hybrid males (p=0.70) that pupated on the food, while F1 hybrid females pupated on the food significantly less often compared to simX hybrid males (p<0.0001 after correcting for multiple comparisons; Fig 3). The fact that F1 hybrid females behave identically to melX hybrid males indicates that the variation in pupation behavior on the *D. melanogaster* X chromosome (i.e. fewer pupae on the food) is dominant to the pupation behavior on the *D. simulans* X chromosome (i.e. more pupae on the food), because F1 hybrid females have one X chromosome from each species.

The fact that hybrid males and females with a *D. melanogaster* X chromosome behave indistinguishably from their *D. melanogaster* parent strain, and hybrid males with a *D. simulans* X chromosome behave indistinguishably from their *D. simulans* parent strain (Fig 3), suggests that hybrid pupation site choice behavior falls well within the typical range exhibited by either parent strain. We found a similar pattern when we compared the average pupation height of F1 hybrids and their parents: F1 females and melX males did not have significantly different pupation heights compared to their *D. melanogaster* parents, and simX males did not differ from their *D. simulans* parental strain (Fig S3, Table S5). This is important to note, as it implies that hybrids, particularly simX hybrids, are not just unable to climb the vial walls due to developmental inconsistencies caused by intrinsic incompatibilities. If anything, simX hybrids displayed slightly higher proportions of individuals climbing the vial walls (and pupated slightly further from the food) compared to their parent strain, underscoring their vigor.

### 3. A deficiency screen of the X chromosome identifies two regions of interest

We found significant variation in pupation index among the deficiency hybrid crosses (Kruskal-Wallis Test: χ^2^=336.90, df=89, p<0.0001; Fig 4A). A pupation index greater than 1 indicates that more deficiency hybrids pupated on the food than balancer hybrids. This suggests that the *D. melanogaster* deficiency region may be revealing recessive *D. simulans* genetic variation that causes the deficiency hybrids to pupate on the food surface. However, the average pupation index across all 90 deficiency hybrid crosses was 0.88 (Fig 4A), which was significantly lower that our expected mean of 1 (Wilcoxon test: p<0.0001). As a result, we compared the pupation index for all deficiencies to our expected value of 1 and to the grand mean pupation index for these lines (0.88). Six deficiencies had pupation indices significantly greater than one: (Df(1)ED411, Df(1)BSC869, Df(1)ED6720, Df(1)ED6906, Df(1)BSC530, and Df(1)Exel6255). Three of these, (Df(1)BSC869, Df(1)ED6906, and Df(1)Exel6255), remained significant after sequential Bonferroni correction for multiple comparisons (Fig 4A; Table S2). Because the other 3 deficiencies, (Df(1)ED411, Df(1)ED6720, and Df(1)BSC530), had pupation indices significantly greater than 0.88 after sequential Bonferroni correction (Table S2), we included them in our list of potential deficiencies of interest.

**Fig 4.**
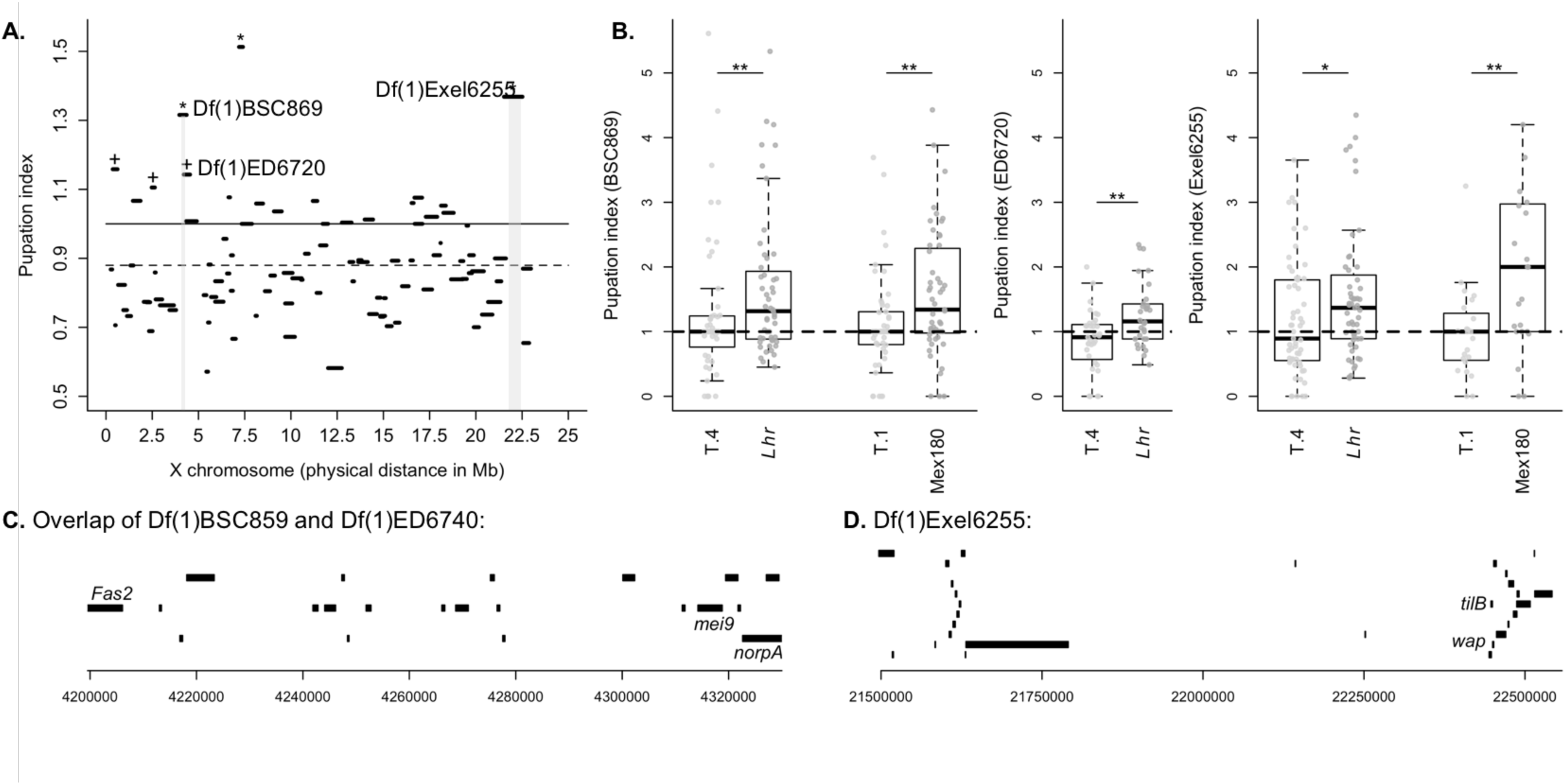
Hybrid deficiency screen of the X chromosome identifies two regions of interest. **A.** The median pupation index for each of the 90 deficiency hybrid crosses (y-axis) is plotted by the physical map distance each engineered deletion spans along the X chromosome (x-axis). Deficiencies with a pupation index significantly greater than 1 (solid line) after correction for multiple comparisons are denoted by an asterisk (significance levels for all deficiencies are listed in Table S2). Deficiencies with a median pupation index significantly greater than 0.88 (the grand mean, dashed line) after correction for multiple comparisons are denoted with a cross. All of these lines also had pupation indices significantly greater than 1 before correcting for multiple comparisons. The two regions we pursued for candidate gene validation are highlighted in light grey. Note, we did not pursue the remaining three significant deficiency strains because they showed similar pupation indices when crossed to *D. melanogaster* (Fig S4). **B.** The pupation indices of the deficiencies from the grey highlighted areas in Part A are shown for the original hybrid cross (*Lhr*) and for a cross to the T.4 *D. melanogaster* strain. BSC869 and Exel6255 were additionally crossed to *D. simulans* strain Mex180 and *D. melanogaster* strain T.1. Asterisks denote a significant difference between pupation indices of deficiency strains crossed to *D. melanogaster* and *D. simulans* (* = p < 0.05, ** = p < 0.01). **C.** The region uncovered by the overlap of deficiencies BSC869 and ED6720 and the 23 genes contained within it. Direction of the gene transcript is denoted with an arrow. The three genes with available disruption strains *(Fas2*, *mei9*, and *norpA*) are labelled. **D.** The region uncovered by Exel6255 and the 28 genes contained within it. Direction of the gene transcript is denoted with an arrow. The two genes with available disruption strains *(tilB* and *wap*) are labelled.

To ensure that the deficient region actually reveals *D. simulans* variation contributing to pupation site choice behavior, rather than creating lines that behave abnormally due to the extended hemizygosity within the deficiency region, we crossed each of the six significant deficiencies listed above to the T.4 wild-type *D. melanogaster* strain. For two of the six deficiency strains, Df(1)ED411 and Df(1)BSC530, we found no difference in the pupation index when crossed to *D. melanogaster* compared to the pupation index when crossed to the *Lhr* strain (Wilcoxon tests; Df(1)ED411: n= 29-36, χ^2^=1.74, p=0.19, Fig S4A; Df(1)BSC530: n=18-22, χ^2^=1.47, p=0.23, Fig S4C), and Df(1)ED6906 had a significantly higher pupation index when crossed to *D. melanogaster* (n= 26-33, χ^2^=9.42, p=0.002; Fig S4B). These results suggest that the phenotypes of these three lines are a result of their hemizygosity within the deficiency region, as the deficient *D. melanogaster* females still pupate on the food more often than the balancer females. Note, this effect is unlikely to be driven by the balancer females, as the same balancers occur in many of the other deficiency lines we screened.

When we crossed the remaining three deficiency strains, Df(1)BSC869, Df(1)ED6720, and Df(1)Exel6255, to the T.4 *D. melanogaster* strain, we found a pupation index significantly lower than the index we calculated when crossing to *Lhr* (Wilcoxon tests; Df(1)BSC869: n= 51, χ^2^=4.35, p=0.037; Df(1)BSC6720: n= 30-35, χ^2^=7.90, p=0.0049; Df(1)Exel6255: n= 55-57, χ^2^=4.68, p=0.0306; Fig 4B), and no different than one (Df(1)BSC869: p=0.46; Df(1)BSC6720: p=0.10; Df(1)Exel6255: p=0.50). This result suggests that these deficient regions do reveal recessive *D. simulans* variation affecting pupation site choice behavior in hybrids, but have no effect when made hemizygous in *D. melanogaster*. Two of these three deficiencies overlap: Df(1)BSC869 and Df(1)ED6720 (Fig 4A). To further confirm that this pattern is not unique to these specific *D. melanogaster* and *D. simulans* wild-type strains, we crossed one of the overlapping deficiency strains, Df(1)BSC869, and Df(1)Exel6255 to another *D. melanogaster* wild-type strain (T.1) and another *D. simulans* wild-type strain (Mex180). The pattern remained consistent for both of these deficiencies: when crossed to *D. melanogaster*, the pupation index was significantly lower than when crossed to *D. simulans* (Fig 4B; Wilcoxon tests: Df(1)BSC869: n=50-51, χ^2^=7.78, p=0.0053; Df(1)Exel6255: n= 23, χ^2^=7.74, p=0.0054).

One deficiency strain, Df(1)ED7265, had a pupation index significantly lower than the grand mean of all the lines (0.88) following Bonferroni correction for multiple comparisons (Table S2). This suggests the potential for a region of the *D. simulans* X chromosome with transgressive effects—that is, a locus that causes *D. simulans* larvae to pupate farther from the food surface. While this is certainly interesting, we did not pursue this region because its effect is contrary to the species-wide difference we found. This locus may be interesting for further investigation, however, as it could contribute to the significant variation in pupation behavior we observed among *D. simulans* strains (Fig 2).

### 4. Gene knockouts and RNAi knockdown suggest that *Fas2* is involved in divergent pupation behavior

We tested knockouts of three candidate genes in our first region of interest: the overlap of Df(1)BSC869 and DF(1)ED6720, excluding the region covered by DF(1)ED6727 (Fig 4C). We found no significant difference in the pupation index obtained when we crossed the *mei-9^A1^* mutant allele to the *Lhr D. simulans* strain and the T.4 *D. melanogaster* strain (Wilcoxon test: n= 51-52, χ^2^=0.37, p=0.54; Fig S5A), indicating that *mei9* is unlikely to be involved in pupation site choice. Similarly, we found no significant differences in the pupation behavior of female hybrids containing the *norpA^36^* mutant allele and female hybrids containing a *norpA* rescue allele (Wilcoxon test: n= 22-24, χ^2^=3.10, p=0.08 Fig S6), with the knockout hybrids actually having a slightly lower proportion of flies pupating on the food. These data suggest that gene knockouts with expression in the larval nervous system do not, in general, increase the number of hybrids pupating on the food.

In contrast, when we crossed the mutant allele *Fas2^eb112^* to *D. simulans* (*Lhr*) and *D. melanogaster* (T.4), we found that *Fas2^eb112^* hybrids had a significantly higher pupation index than the *D. melanogaster* knockouts (Wilcoxon test: n= 50-51, χ^2^= 6.97, p= 0.0083; Fig 5A), suggesting that *Fas2* may be involved in pupation site choice. To ensure this pattern is not unique to these strains, we crossed *Fas2^eb112^* to additional *D. simulans* (Mex180) and *D. melanogaster* (T.1) wild-type strains. We again found the same pattern: the pupation index for knockout hybrids was significantly higher than for *D. melanogaster* knockouts (Wilcoxon test: n= 52-53, χ^2^= 27.2, p<0.0001; Fig 5A). As further verification, we tested a second *Fas2* strain: a p-element insertion allele, *Fas2^G0293^*, and similarly found that the pupation index for *Fas2^G0293^* hybrids (crossed to *Lhr*) was significantly higher than that for *D. melanogaster* knockouts (crossed to T.4; Wilcoxon test: n= 52-53, χ^2^= 9.73, p=0.0018; Fig 5B). Although the nature of the lesion is uncharacterized for *Fas2^G0293^*, these results suggest that it is indeed a loss of function allele, as it behaves indistinguishably from the verified knockout *Fas2^eb112^*. We used the consensus combined p-value test (Rice, 1990), which tests the combined effect of independent tests of the same hypothesis, to look at the overall pattern for *Fas2^eb112^*, and *Fas2* as a whole (i.e. including results from both *Fas2^eb112^* and *Fas2^G0293^*), and found a strongly significant pattern of higher pupation indices for hybrid crosses compared to *D. melanogaster* crosses (*Fas2^eb112^*: p= 3.59 × 10^−6^; *Fas2*: p= 2.35 × 10^−8^).

**Fig 5.**
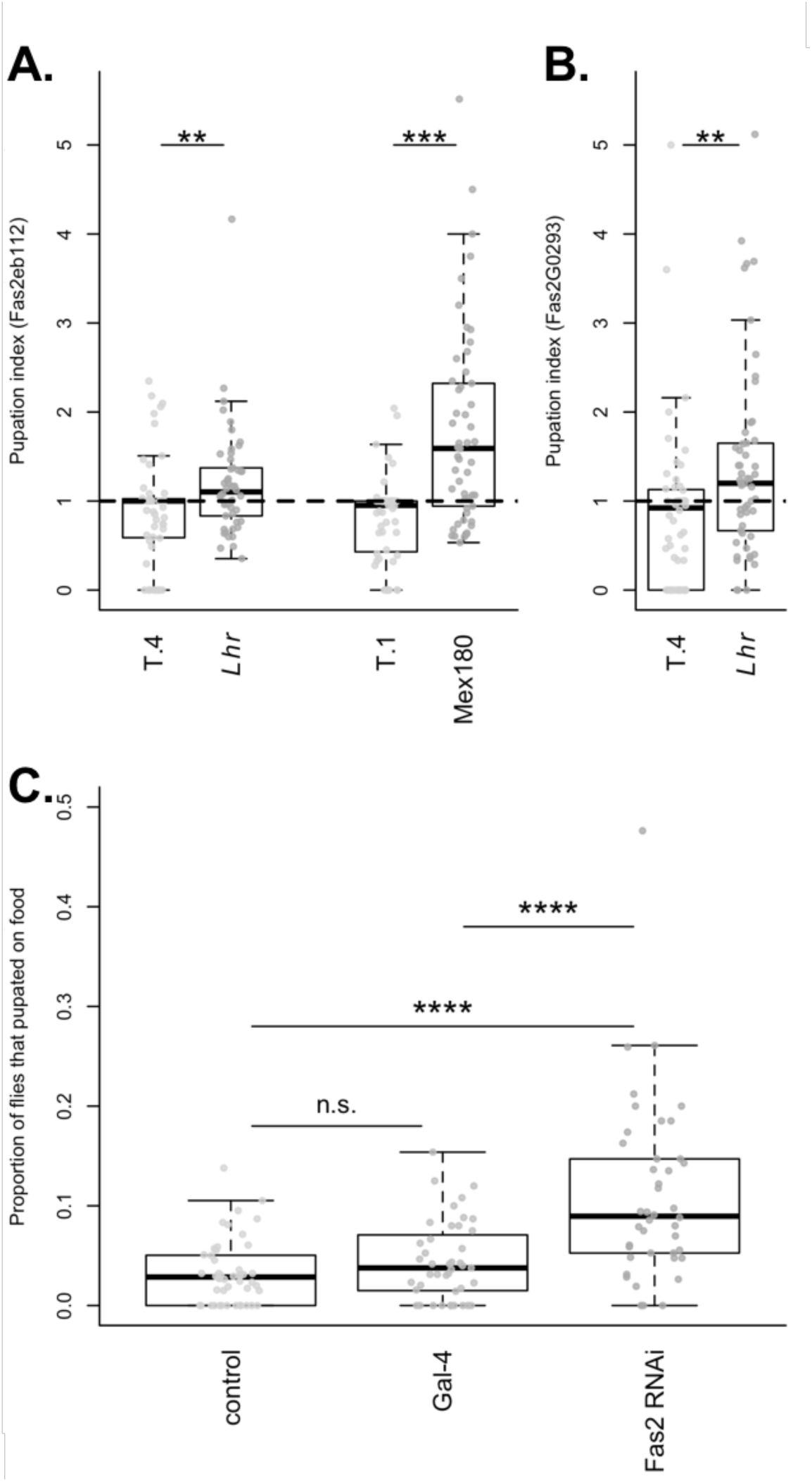
Knockouts and RNAi knockdown confirm the role of *Fas2* in evolved differences in pupation site choice. **A.** The pupation indices of the *Fas2^eb112^* gene disruption are shown for the comparison between the original hybrid cross (*Lhr*) and a cross to the *D. melanogaster* T.4 strain. Also shown is the comparison for crosses to the *D. simulans* strain Mex180 and the *D. melanogaster* strain T.1. **B.** The pupation indices for a second gene disruption, *Fas2^G0293^*, are shown for the original hybrid cross (*Lhr*) and a *D. melanogaster* strain (T.4). For A and B, asterisks denote significance (** = p < 0.01, **** = p<0.0001; N = 50-53). **C.** The results of pan-neuronal knockdown of the *Fas2* transcript via RNAi. The proportion of RNAi/control individuals found on the food are shown for the control cross (elav-Gal4 driver crossed to the RNAi background stock), the Gal-4 hairpin RNA cross (Gal-4), and *Fas2* RNAi cross. For C, asterisks denote significance after correcting for multiple comparisons (**** = p<0.0001; N = 42-48).

Next, we used RNAi with the elav-Gal4 driver to reduce expression of *Fas2* throughout the nervous system in *D. melanogaster*. We found that a significantly higher proportion of *Fas2* RNAi flies pupated on the food compared to the control flies from either the background (Wilcoxon test: n= 42-48, p<0.0001 after sequential Bonferroni correction) or Gal4-1 cross (Wilcoxon test: n= 42-43, p<0.0001 after sequential Bonferroni correction; Fig 5C); these results are unchanged when we control for density effects (Fig S7A). Similarly, when we compared pupation height, we found that RNAi flies pupated significantly closer to the food compared to both the background (Wilcoxon test: n=42-48, p<0.0001 after sequential Bonferroni correction) and Gal-4 crosses (Wilcoxon test: n= 42-43, p<0.0001 after sequential Bonferroni correction; Fig S8A), providing further evidence for *Fas2*’s role in pupation site choice.

### 5. Gene knockouts and RNAi knockdown suggest that *tilB* is involved in divergent pupation behavior

We tested knockouts of two candidate genes in our second region of interest: DF(1)Exel6255 (Fig 4D). We found no significant difference in the pupation index obtained when we crossed the *wap^2^* mutant allele to the *Lhr D. simulans* strain and the T.4 *D. melanogaster* strain (Wilcoxon test: n= 51-55, χ^2^=0.32, p=0.57; Fig S5B), indicating that *wap* is unlikely to be involved in pupation site choice. In contrast, when we crossed the *tilB^1^* and *tilB^2^* mutant alleles to *D. simulans* (*Lhr*) and *D. melanogaster* (T.4), we found that the *tilB* knockout hybrids had significantly higher pupation indices than the *D. melanogaster* knockouts for both alleles (Wilcoxon tests; *tilB^1^*: n= 56, χ^2^= 6.61, p= 0.0101; Fig 6A; *tilB^2^*: n= 57, χ^2^= 6.61, p= 0.0101; Fig 6B). To test whether this difference is consistent in other backgrounds, we crossed both the *tilB^1^* and *tilB^2^* mutant alleles to additional *D. simulans* (Mex180) and *D. melanogaster* (T.1) wild-type strains. We had difficulty crossing our *tilB* strains to the Mex180 strain, so our sample sizes for these crosses are smaller, but there is a nonsignificant trend towards a higher pupation index for the knockout hybrids compared to the *D. melanogaster* hybrids for both alleles (Wilcoxon tests; *tilB^1^*: n=36-37, χ^2^= 2.80, p= 0.0943, Fig 6A; *tilB^2^*: n=12-18, χ^2^= 3.44, p= 0.0638, Fig 6B). We used the combined consensus p-value test (Rice, 1990) to look at the overall pattern for *tilB^1^*, *tilB^2^*, and *tilB* as a whole (i.e. including results from both *tilB^1^* and *tilB^2^*), and found a strongly significant pattern of higher pupation indices for hybrid crosses compared to *D. melanogaster* crosses (*tilB^1^*: p= 0.0022; *tilB^2^*: p= 0.0037; *tilB*: p= 2.47 × 10^−5^).

**Fig 6.**
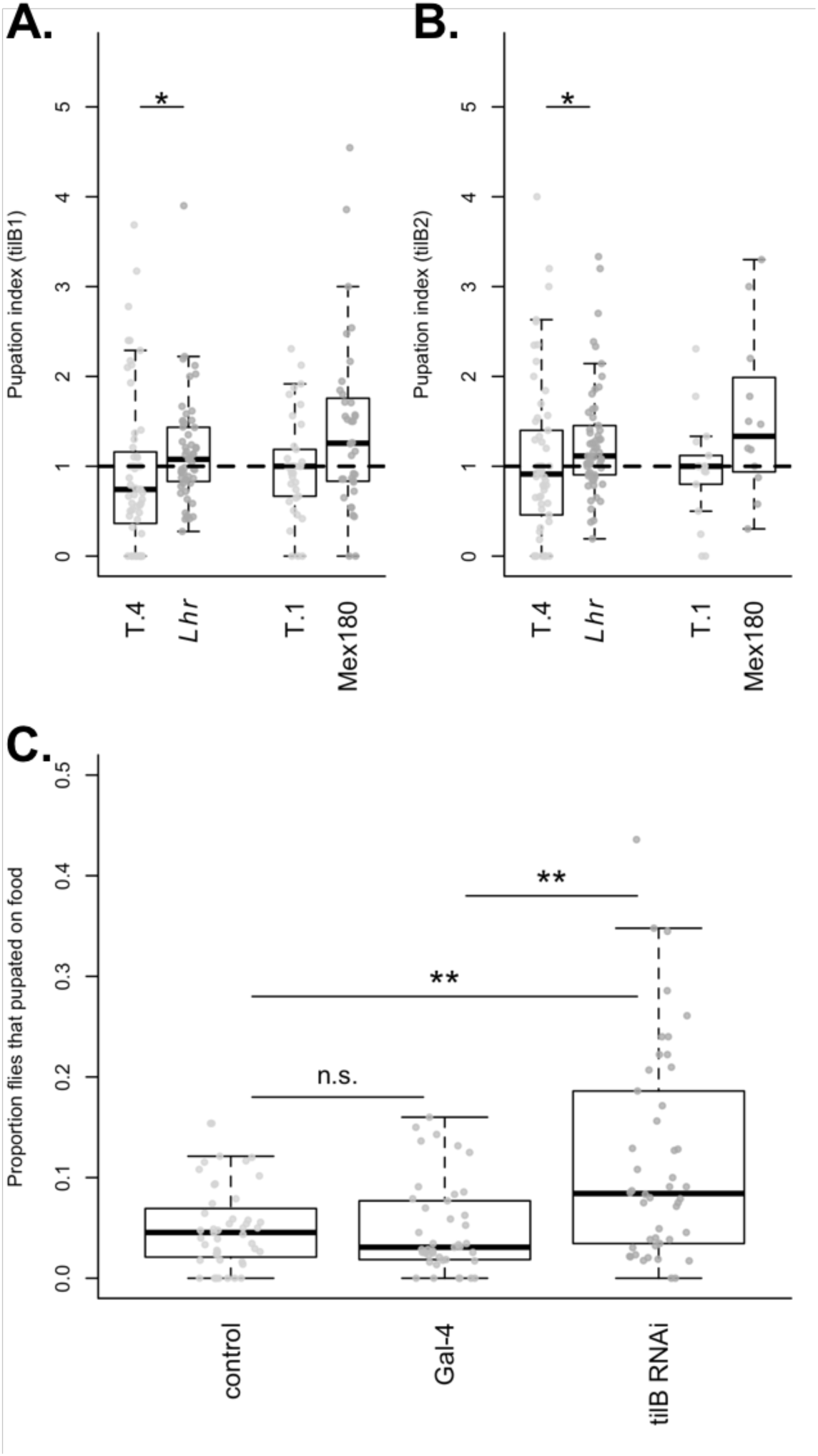
Knockouts and RNAi knockdown confirm the role of *tilB* in evolved differences in pupation site choice. **A.** The pupation indices of the *tilB^1^* gene disruption are shown for the comparison between the original hybrid cross (*Lhr*) and a cross to the *D. melanogaster* T.4 strain. Also shown is the comparison for crosses to the *D. simulans* strain Mex180 and the *D. melanogaster* strain T.1. **B.** The pupation indices for a second gene disruption, *tilB^2^*, are shown for the original hybrid cross (*Lhr*) and the cross to *D. melanogaster* strain T.4, and for crosses to the *D. simulans* strain Mex180, and the *D. melanogaster* strain T.1. For A and B, asterisks denote significance (* = p < 0.05, ** = p = 0.01; N = 41-47). **C.** The results of pan-neuronal knockdown of the *tilB* transcript via RNAi. The proportion of RNAi/control individuals found on the food are shown for the control cross (elav-Gal4 driver crossed to the RNAi background stock), the Gal-4 hairpin RNA cross (Gal-4), and *tilB* RNAi cross. For C, asterisks denote significance after correcting for multiple comparisons (* = p < 0.05, ** = p < 0.01; N = 41-47).

As for *Fas2* above, we then used RNAi with the elav-Gal4 driver to reduce expression of *tilB* throughout the nervous system in *D. melanogaster*. We found that a significantly higher proportion of RNAi flies pupated on the food compared to the control flies from either the background (Wilcoxon test: n= 46-47, p<0.01 after sequential Bonferroni correction) or Gal4-1 cross (Wilcoxon test: n= 41-46, p<0.01 after sequential Bonferroni correction; Fig 6C); these findings are consistent when we control for density effects (Fig S7B). In addition, when we compared pupation height, we found that RNAi flies pupated significantly closer to the food compared to both the background (Wilcoxon test: n=46-47, p<0.0001 after sequential Bonferroni correction) and Gal-4 crosses (Wilcoxon test: n= 46-47, p<0.0001 after sequential Bonferroni correction; Fig S8B), providing additional support for *tilB*’s role in pupation site choice.

### 6. *tilB* is expressed more highly in *D. melanogaster* strains

We performed qRT-PCR to quantify relative *tilB* transcript abundance for two strains of *D. simulans* (Per005 and Geo288) and two strains of *D. melanogaster* (CA1 and T.4). For the two *D. simulans* strains and the Geo288 *D. melanogaster* strain, we collected larvae from two stages of larval development (96 and 120 hours following oviposition). For the other *D. melanogaster* strain (T.4), we were only able to obtain enough larval tissue at 96 hours following oviposition, due to low fecundity. Because the relative transcript abundance data had a skewed distribution, we used the reciprocal root transformation to normalize the species and overall distributions (Shapiro-Wilk tests for normality: all p >0.34). We then analyzed the relative transcript abundance of *tilB* using a nested ANOVA with the following factors: species, strain nested within species, larval age (96 and 120 hours), and the interaction between species and larval age. The interaction term between species and larval age was not significant (p=0.41), so we removed it from the model. We found that larvae from the 120-hour sampling period had significantly lower *tilB* expression than 96-hour larvae (F_1,14_= 6.16, p= 0.026). While we did not detect any significant differences between the 2 strains from the same species (F_2,14_=2.39, p=0.13), we found a significantly higher average relative amount of *tilB* transcript in *D. melanogaster* larvae compared to *D. simulans* larvae (F_1,14_= 9.74, p= 0.0075; Fig 7).

**Fig 7.**
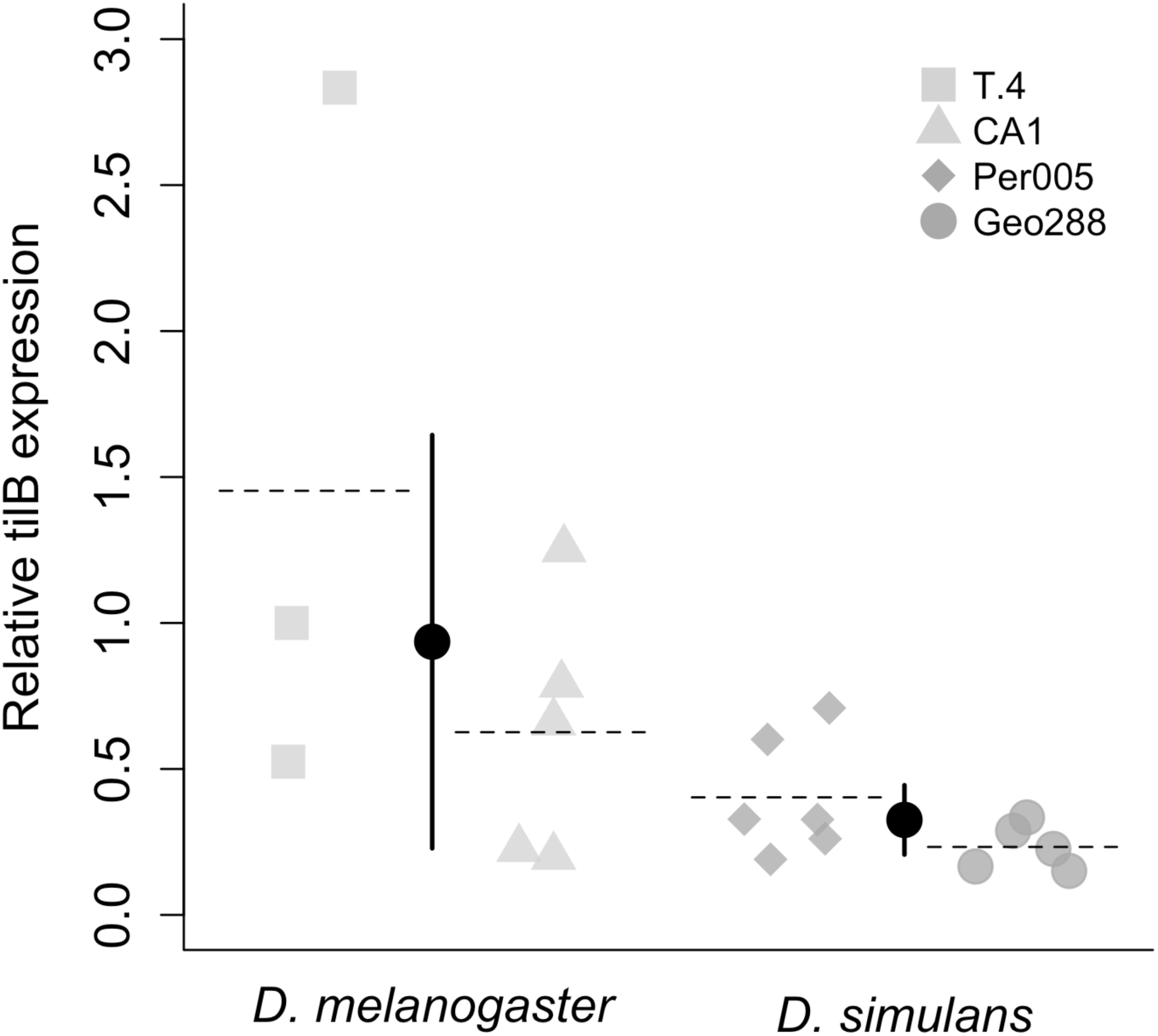
Relative differences in *tilB* transcript abundance between species. The relative abundance of *tilB* transcript detected by qRT-PCR in *D. melanogaster* and *D. simulans*. Each data point represents the average of two technical replicates for a single biological replicate. Relative transcript abundance is significantly higher in *D. melanogaster* strains on average (p < 0.01). Light grey squares (T.4) and triangles (CA1) represent the two *D. melanogaster* strains measured, while dark grey diamonds (Per005) and circles (Geo288) represent the two *D. simulans* strains. The dashed lines depict the average expression across both time points for each strain. The black circles represent the species-wide mean, and error bars depict the 95% confidence interval surrounding the mean. Note that the exact rank of relative *tilB* expression is perfectly inversely correlated to the pupation indices of each strain (i.e., the strain with the highest *tilB* expression has the lowest proportion of pupae on the food, and so on).

Because we performed qRT-PCR on an extreme strain and a strain closer to the average for each species, these four lines represent a continuum of pupation site choice behavior, with T.4 (*D. melanogaster*) having the lowest proportion of pupae on the food, followed by CA1 (*D. melanogaster*), then Per005 (*D. simulans*), and last, Geo288 (*D. simulans*) having the highest proportion of pupae on the food (Fig 2). These four strains follow an identical pattern for *tilB* gene expression, with T.4 having the highest relative transcript abundance, and Geo288 having the lowest (Fig 7). Although it is not possible to detect a significant effect with a sample size of 4, this suggests that *tilB* gene expression may be negatively correlated with the proportion of pupae on the food (Spearman’s rank correlation: r_s_ = −1, p< 0.10).

### 7. Effect sizes

We used the results above to estimate how much of the difference in pupation site preference between *D. melanogaster* and *D. simulans* can be attributed to the X chromosome, and found that the X accounts for approximately 55.6% (95% CI= 31.4%-80.2%) of the difference. We then estimated the effect size of our deficiencies of interest, and found that the overlap of Df(1)BSC869 and DF(1)ED6720 explains approximately 44.1% (95% CI= 33.5%-74/5%) of the X chromosome effect, and Df(1)Exel6255 explains approximately 52.3% (95% CI= 33.7%-85.9%) of the X effect. Finally, we estimated the effect size of our two identified candidate genes, *Fas2* and *tilB*. We found that *Fas2* explains approximately 93% (95% CI= 78.5%-134.9%) of the pupation difference attributed to the overlap between Df(1)BSC869 and DF(1)ED6720, and 41% (95% CI= 33.8%-76.5%) of the pupation difference attributed to the X chromosome. Similarly, we found the *tilB* explains approximately 87.2% (95% CI= 66.2%-135.9%) of the pupation difference attributed to Df(1)Exel6255, and 45.6% (95% CI= 35.5%-76.1%) of the pupation difference attributed to the X chromosome.

It is important to note that our effect size estimate for the X chromosome may be an overestimate, as calculating effect sizes using only reciprocal hybrids does not account for potential transgressive autosomal effects (Mittleman et al., 2017). While subject to the same potential effects, the effect size of the deficiencies and individual gene knockouts were calculated as proportions of the total X effect (and also of the deficiency effect for gene knockouts), and thus represent a more accurate estimate of their overall contribution, regardless of the total X effect. Still, these estimates could be affected by variation in the different *D. melanogaster* backgrounds in which the deficiencies and knockout strains were made.

## Discussion

### A species-level difference in pupation site choice behavior

Our initial survey of pupation site choice behavior in *D. melanogaster* and *D. simulans* expands upon a previously reported interspecific difference (Markow, 1979). Consistent with these previous results, on average, *D. simulans* strains had a greater proportion of flies pupating on the food surface. However, we used 11 *D. melanogaster* and 12 *D. simulans* strains sourced from around the globe to demonstrate this difference. While the species difference holds when comparing the grand mean of all strains for each species, there is substantial variation within species. This variation is so significant that the species’ distributions overlap, with some *D. simulans* strains, like Mex180 and Cal006, more closely resembling *D. melanogaster* strains in pupation behavior (Fig 2). These documented differences in pupation behavior among *Drosophila* species, in combination with our understanding of the environmental variables that affect this behavior within species (Hodge & Caslaw, 1998; Sameoto & Miller, 1968b; Seyahooei et al., 2009; Sokal et al., 1960), will be useful in identifying the selection pressures (if any) that affect the evolution of this trait. Differences in pupation site choice behavior may be a form a niche partitioning where species co-occur, as has been previously suggested (Arthur & Middlecote, 1984). Alternatively, pupation site choice may be an adaptive response to parasite or parasitoid presence (Kraaijeveld & Godfray, 2003). A globally sourced panel of lines with significant variation, such as we describe, provides an inroad for studies comparing pupation behavior to differences in the ecology of each collection site, such that we can better understand the ultimate causes of this behavioral evolution.

### The genetic architecture of pupation site choice behavioral evolution

Using hybrid crosses in the same background that controlled for maternal inheritance, we were able to estimate the effect of the X chromosome on pupation behavior. We found no evidence that hybrid behavior was atypical relative to either parent strain, suggesting these measurements are indeed reliable. We found a significant effect of the X, in that reciprocal hybrid males pupate in similar locations as the X-donating parent. We calculate that this chromosome explains 55.6% (95% CI= 31.4%-80.2%) of the total phenotypic difference between parent strains. Although the exact contribution of the X chromosome reported here may be an overestimate due to transgressive autosomal effects that cannot be detected in a hybrid background (Mittleman et al., 2017), a similar X-effect has been detected for pupation behavior when comparing *D. simulans* and *D. sechellia* (Erezyilmaz & Stern, 2013).

We also found that hybrid females, which inherit one X chromosome from each parent, pupate like the *D. melanogaster* parent strain. This suggests that *D. melanogaster* alleles are dominant to *D. simulans* alleles. The dominance of *D. melanogaster* alleles makes it possible to use engineered deletions, available in *D. melanogaster* strains, to map regions containing recessive *D. simulans* variation affecting pupation behavior (Cook et al., 2012).

The three significant deficiencies identify two regions of the X chromosome with detectable effects on pupation behavior: one spans X:4,204,351 – 4,325,174 and explains ∼44.1% of the X-effect, and the other spans X:21,519,203 – 22,517,665 and explains ∼52.3% of the X-effect. These regions contain 23 and 28 genes, respectively. Our analysis of the available gene knockouts within these regions points to two genes: *tilB* and *Fas2* (see below). We calculate that *tilB* and *Fas2* explain the majority of the effect of their respective deficiency regions (point estimate = 87.2%, 95% CI= 66.2%-135.9%; point estimate = 93%, 95% CI= 78.5%-134.9%, respectively). Taken together, our results suggest that a substantial share of the difference in pupation behavior between *D. simulans* and *D. melanogaster* can be attributed to just two genes.

### *tilB* and *Fas2*: loci of evolution for divergent pupation behavior between *D. simulans* and *D. melanogaster*

We have presented substantial evidence for a role of both *tilB* (*touch insensitive larva B*) and *Fas2* (*Fasciclin* 2) in the divergence of pupation behavior among these species. For each locus, we have shown that two independent knockouts replicate the pattern of the regions identified by the deficiency screen: hybrid females hemizygous for the *D. simulans* locus pupate on the food surface significantly more often than females with both the *D. simulans* and *D. melanogaster* loci.

We then used RNAi knockdown of each gene in *D. melanogaster* to show that reduced expression of *tilB* and *Fas2* transcripts results in a more *D. simulans*-like pupation site choice behavior. To directly test for differences in expression of *tilB*, we performed qRT-PCR during two larval developmental time points. Congruent with our RNAi knockdown results, we find that mean relative transcript abundance is significantly higher in *D. melanogaster* larvae than in *D. simulans* larvae. Interestingly, we surveyed two lines per species, representing a continuum of pupation site choice behavior, and found that the mean proportion of pupae on the food surface for these lines was perfectly negatively correlated with *tilB* gene expression. Although these results are only for 4 lines, they suggest that lower *tilB* gene expression may be associated with a higher proportion of larvae pupating on the food surface.

While our present study does not present a functional analysis of the *D. melanogaster* or *D. simulans Fas2* or *tilB* alleles, we can use the *D. melanogaster* annotation of each gene to speculate about their role in the evolution of pupation behavior. *Fas2* is a large gene, spanning over 70,000 base pairs, with expression peaking during the larval wandering stage (L3) (Graveley et al., 2011). It is also complex, with seven transcripts composed of various combinations of 16 exons. Broadly, *Fas2* functions as a neuronal recognition molecule, and is involved in patterning the larval nervous system (Grenningloh et al., 1991). Expression of *Fas2* is critical for synapse formation and growth at the larval neuromuscular junction (Davis et al., 1997; Schuster et al., 1996) and is also important for patterning of the larval mushroom body (Kurusu et al., 2002). With expression in both the central and peripheral nervous system, it is possible that differences at the *Fas2* locus differentially wire the *D. melanogaster* and *D. simulans* brains, altering how larvae perceive or interpret stimuli. Whether these differences are a result of evolution of the protein sequence, and/or spatial or temporal differences in transcript expression remains to be determined.

Unlike *Fas2*, *tilB* is a short gene, spanning just over 1,700 base pairs, with a single transcript composed of 5 exons. *tilB* is also expressed in wandering larvae and pupae, though it shows higher expression in testes of adult males, due to its role in developing sperm flagella (Graveley et al., 2011). In fact, *tilB* is associated with ciliary motility (Kavlie et al., 2010), and is a part of the mechanosensory transduction machinery (Göpfert & Robert, 2003). Mutant *tilB* larvae display normal locomotor activity, but have a reduced withdrawal response to physical disturbance (Kernan et al., 1994). Consistent with this finding, our data suggest that changes in *tilB* expression could potentially result in differences in peripheral sensory perception between *D. melanogaster* and *D. simulans* larvae, ultimately influencing larval pupation site choice behavior. A more precise functional analysis of *tilB* is necessary to test this hypothesis.

### Standards of evidence and the challenges of interspecific mapping

Above, we have discussed the results of our interspecific deficiency screen of the *Drosophila* X chromosome. This technique has long been employed to map morphological traits, physiological traits, hybrid incompatibility loci, (Barbash et al., 2003; Bour et al., 2000; Cattani & Presgraves, 2012; Cote et al., 1986; Konopka & Benzer, 1971; Pardy et al., 2018; Sawamura et al., 2004), and behaviors (Fanara et al., 2002; Laturney & Moehring, 2012; Moehring & Mackay, 2004). Nonetheless, these screens have been criticized due to their susceptibility to epistatic interactions that can produce false positives (Anholt & Mackay, 2004) and for the imperfect comparison of deficiency chromosomes to balancer chromosomes (Stern, 2014). These problems may be exacerbated in a hybrid background. These issues are potentially reflected by the global average pupation index for these deficiencies being 0.88, rather than 1 (Fig 4A), indicating that, on average, more balancer hybrids pupated on the food compared to deficiency hybrids.

We attempted to exclude the possibility of false positives in our initial deficiency screen in two ways. First, we crossed each deficiency with a significant pupation index to *D. melanogaster* to test for deleterious effects of the deficiencies themselves. Indeed, 3 of 6 significant deficiencies produced similar results when crossed to *D. melanogaster*, and were discarded. Next, to ensure the patterns shown by the remaining three were not due to epistasis among strains, we crossed each to a second *D. simulans* and *D. melanogaster* strain and found consistent results. Taken together, these results suggest that these three deficiencies are likely revealing recessive *D. simulans* variation, rather than background epistatic interactions. It should be noted, however, that species-wide epistasis cannot be conclusively ruled out; nonetheless, such a result is still biologically relevant and meaningful.

The results of our gene disruption tests further enforce our deficiency screen results, again, with multiple controls to limit the possibility of false positives and/or strain-specific epistatic effects. We tested *D. melanogaster* gene disruption strains in a hybrid background for two or three potential candidate genes that are expressed in the larval nervous system from each region identified by our deficiency screen. For each region, only a single candidate gene (*Fas2* and *tilB*) replicated the deficiency effect, suggesting that disrupting larval nervous system genes in a hybrid background does not generally interfere with pupation behavior. Further, we crossed our *tilB* and *Fas2* gene disruption strains to multiple *D. melanogaster* and *D. simulans* strains, and found no evidence of strain-specific epistatic interactions. Finally, we tested two knockouts of each candidate gene to ensure that our results were not specific to the gene aberration.

In addition to our hybrid mapping experiments, we have shown that a reduction in *Fas2* and *tilB* gene expression via RNAi knockdown in a *D. melanogaster* background leads to a more *D. simulans*-like phenotype. This test is performed in a pure *D. melanogaster* background, eliminating the potential for hybrid background epistasis, and the results are consistent with our gene disruption screen. Additionally, these results hold true when calculating both the proportion of flies pupating on the food surface, and the average pupation height. This result is even further corroborated for *tilB*, which shows higher relative expression in *D. melanogaster* compared to *D. simulans* strains (Figure 7). Taken together, our hybrid gene disruption screens and gene expression studies in *D. melanogaster* make an excellent case for a role of *tilB* and *Fas2* in divergent pupation site choice behavior.

Higher standards of evidence have been suggested for mapping in *Drosophila* – the reciprocal hemizygosity test being the gold standard (Stern, 2014). In this study, we performed one half of this test (hybrid females with a disrupted *D. melanogaster* X chromosome and an intact *D. simulans* X chromosome). Unfortunately, the reciprocal half of the test (hybrid females with a disrupted *D. simulans* X chromosome and an intact *D. melanogaster* X chromosome) is not possible in this case for several reasons. The hybrid cross is only successful in one direction (*D. simulans* males crossed to *D. melanogaster* females), so we would need to use *D. simulans* gene disruption males to create the reciprocal cross. This is not an option, however, as *tilB* disruptions are male sterile, and *Fas2* disruptions are male lethal.

A test that could potentially support our findings would be the addition of dominant *D. melanogaster Fas2 and tilB* alleles to a *D. simulans* X background to recover the *D. melanogaster* behavior of pupating off the food surface. We previously performed a similar experiment in males using *D. melanogaster* Y-linked X duplication strains (Table S6) (Cook et al., 2010). Unfortunately, making hybrid males that have two broad segments of the X chromosome created flies that pupated almost entirely on the surface of the food (Fig S9). A similar experiment could be performed in females using third chromosome-linked X duplication strains (Venken et al., 2010), however this is likely to produce a similar effect. Further, these experiments are prone to the same potential epistatic interactions as the deficiencies.

While we have presented multiple lines of evidence, all consistently supporting a role for *tilB* and *Fas2* in divergent pupation site choice behavior, these results must be considered in light of the above caveats. We have taken measures to address these caveats as completely as possible, and present our results with a high standard of evidence. Our study highlights the difficulty of interspecific mapping in producing conclusive results, and underscores a need for transgenic tools to be developed in non-model *Drosophila* species.

### Areas for future research

Our results highlight two main areas for further research. First, while we now have a better understanding of the genetic underpinnings of pupation site choice behavior evolution, it is unclear why these differences evolved in the first place. The variation we recorded among globally sourced *D. melanogaster* and *D. simulans* strains provides a valuable tool with which to pursue this line of inquiry. Second, while we have provided evidence for two specific genes involved in the evolution of this phenotype, we have little understanding of their function in pupation site choice behavior, or how exactly these functions have evolved. The results of our expression study suggest a role of expression differences for *tilB*, but further functional follow-up is necessary to identify the precise molecular underpinnings of pupation site choice preference.

## Acknowledgments

We would like to thank Alexandra Lamoureux, Susanne Tilk, Nastaran Hosseinipour, Sayeh Akhavan, Wesley Cochrane and Veronica Cochrane for their assistance in the collection of pupation preference data. We also thank Daniel Eberl, Stuart Macdonald, Brian McCabe, and William Rice for providing fly stocks used for this study. This work would not have been possible without the community-supported resources available at the Cornell *Drosophila* Stock Center (CSBR 1820594), the Bloomington *Drosophila* stock center (NIH P400D018537), or FlyBase (flybase.org; Gramates et al., 2017). This work was funded by the National Institutes of Health (R01 GM098614).

## Supplementary Figures and Captions

**Fig S1.**
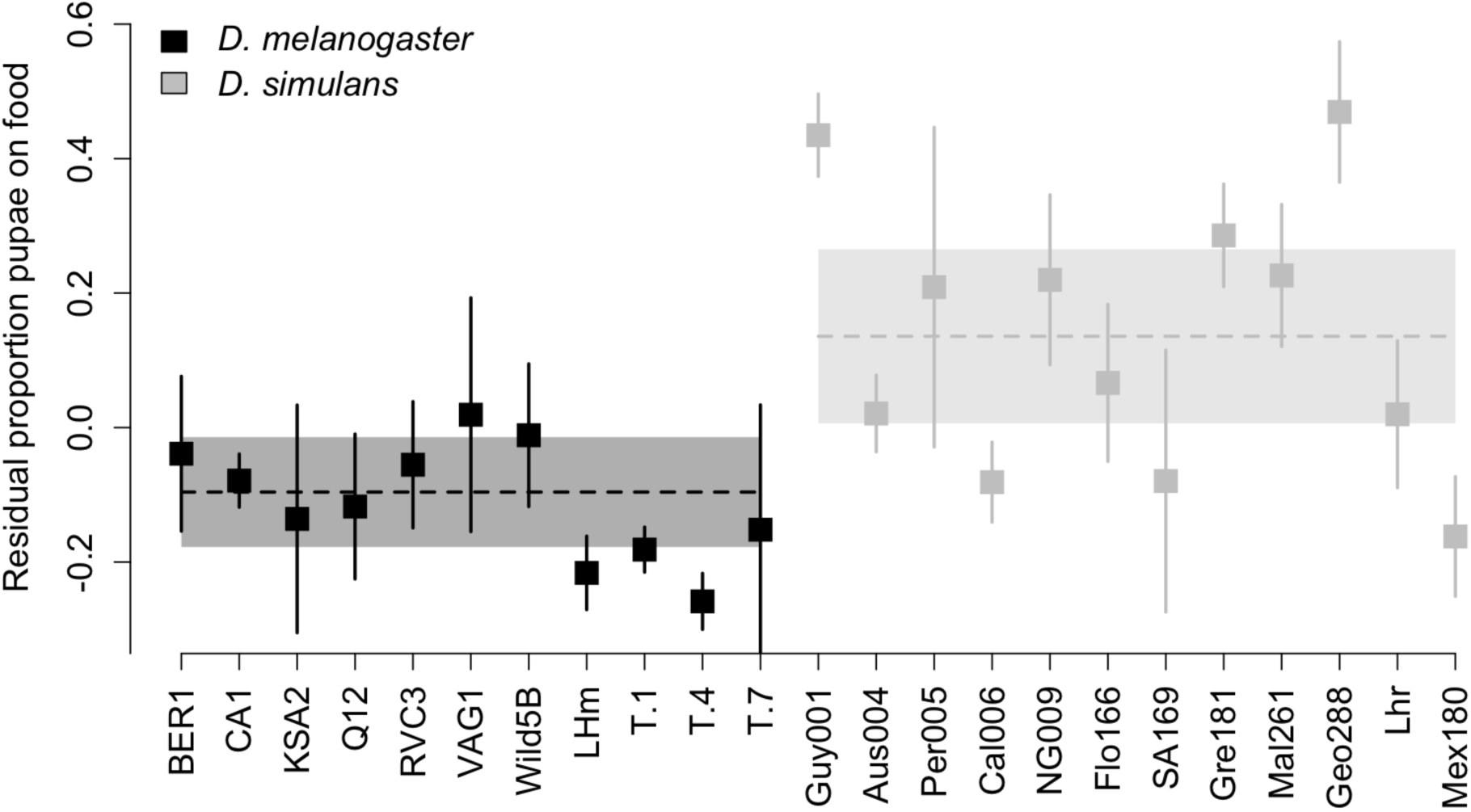
Pupation behavior differences between *D. melanogaster* and *D. simulans* after controlling for density. The mean residuals from a regression between the proportion of pupae on the surface of the food and the total number of pupae in the vial for 11 *D. melanogaster* lines and 12 *D. simulans* lines described in Table S1. Error bars denote the 95% confidence interval around each individual mean (N = 5-8). The dashed horizontal lines indicate the grand mean for each species. The boxes surrounding the dashed lines denote the 95% confidence interval around the grand mean.

**Fig S2.**
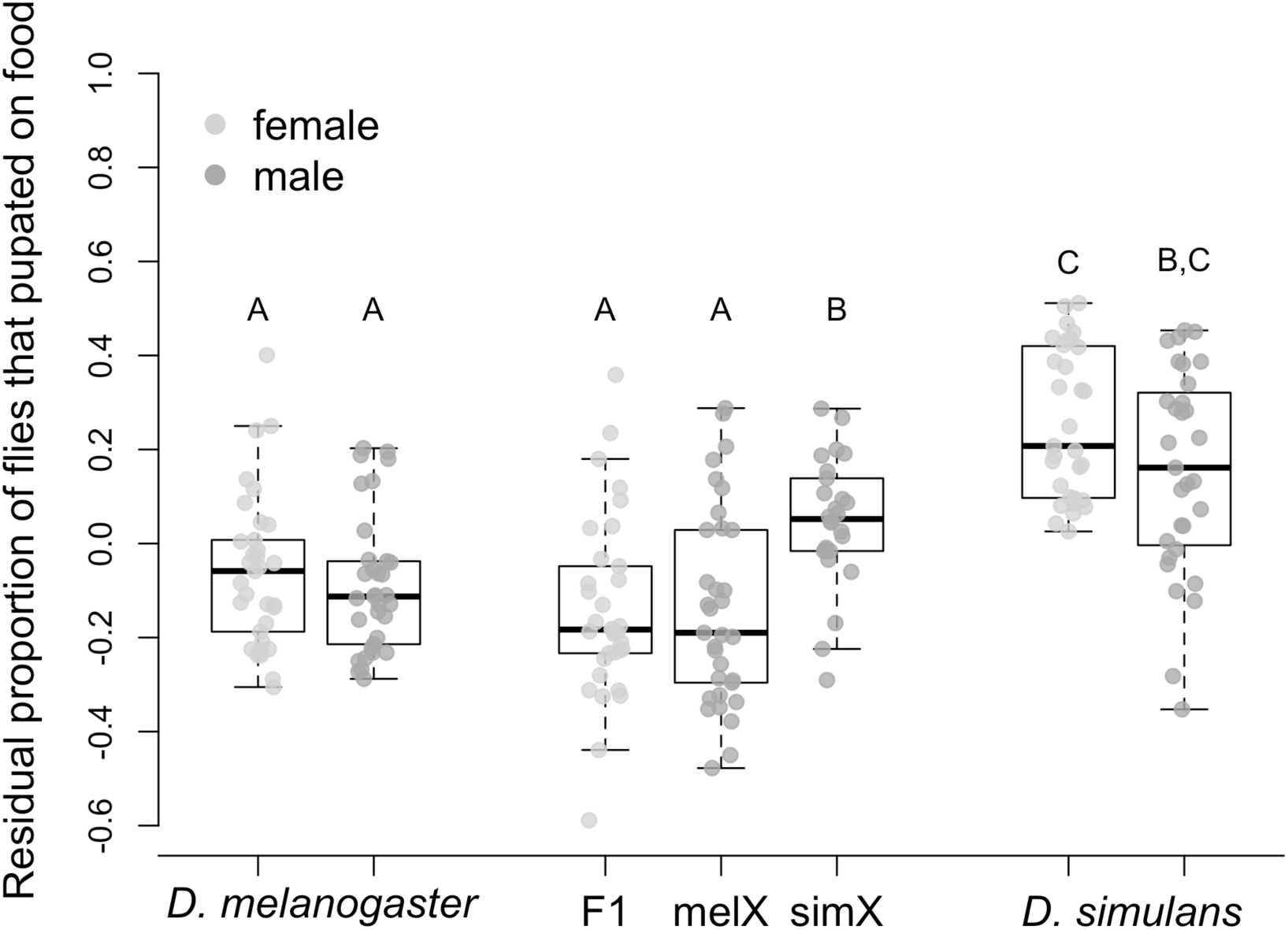
Pupation behavior for *D. melanogaster*, *D. simulans*, and their F1 hybrids after controlling for density. The mean residuals from a regression between the proportion of individuals in a vial that pupated on the surface of the food and the total number of individuals in the vial are shown for males and females from both species and their F1 hybrids. *D. melanogaster* males and females were taken from the LH_M_ strain, while *D. simulans* males and females were taken from the *Lhr* strain. F1 hybrids resulted from a cross between these two strains. The “melX” hybrid males have the *D. melanogaster* X chromosome, and the “simX” hybrid males have the *D. simulans* X chromosome. Both hybrids have *D. melanogaster* cytoplasmic inheritance. Box plots labeled with different letters are significantly different from one another after sequential Bonferroni adjustment for multiple comparisons (p < 0.0001, N = 26-33).

**Fig S3.**
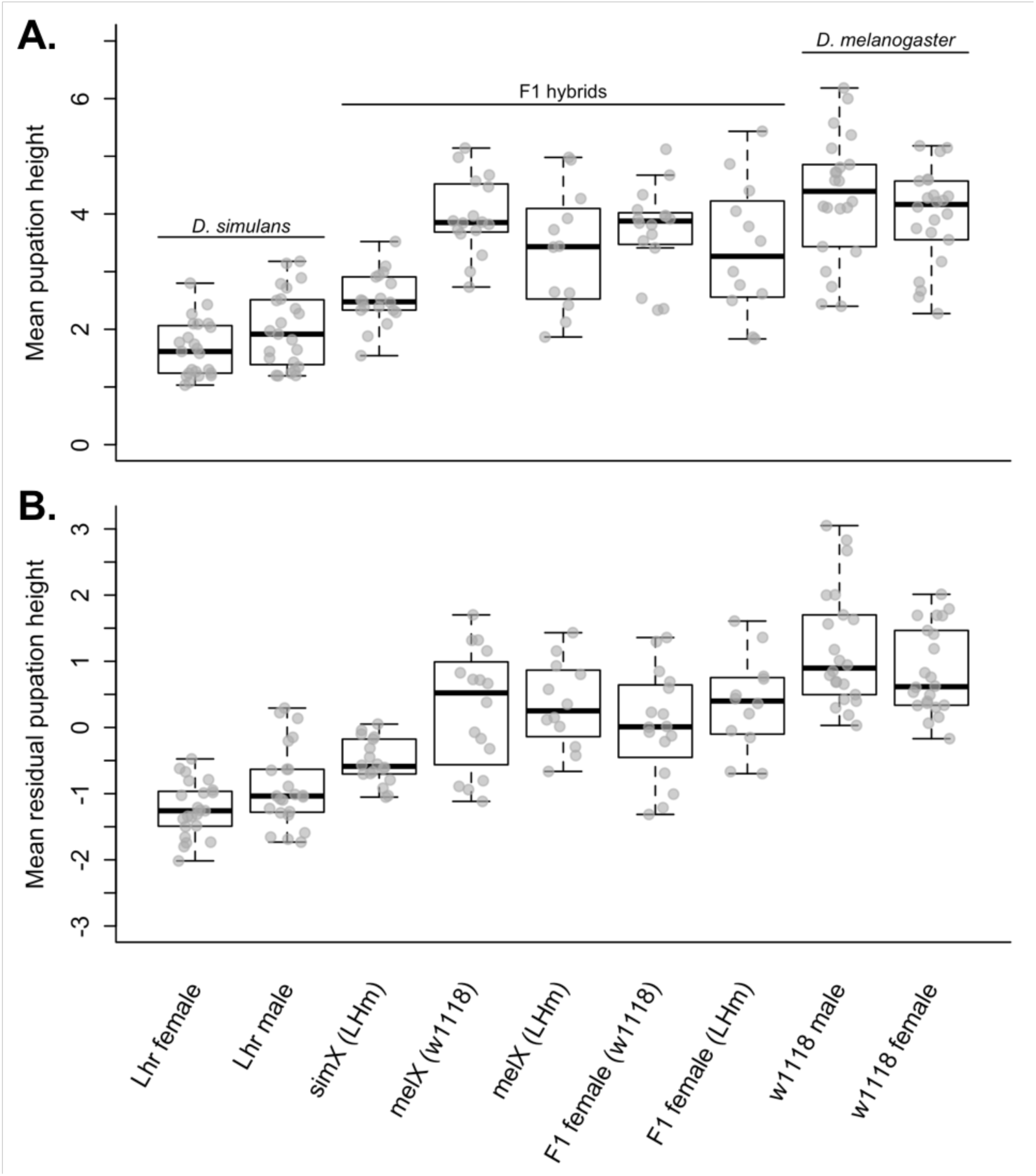
Pupation behavior for *D. melanogaster*, *D. simulans*, and their F1 hybrids, measured as average pupation height within a vial. **A.** The average pupation height is shown for males and females from both species and their F1 hybrids. Higher pupation heights indicate individuals pupated further from the food. **B.** The residuals from a regression between the average pupation height of individuals in a vial and the total number of individuals in that vial are shown for males and females from both species and their F1 hybrids. The results from pairwise statistical tests can be found in Table S5.

**Fig S4.**
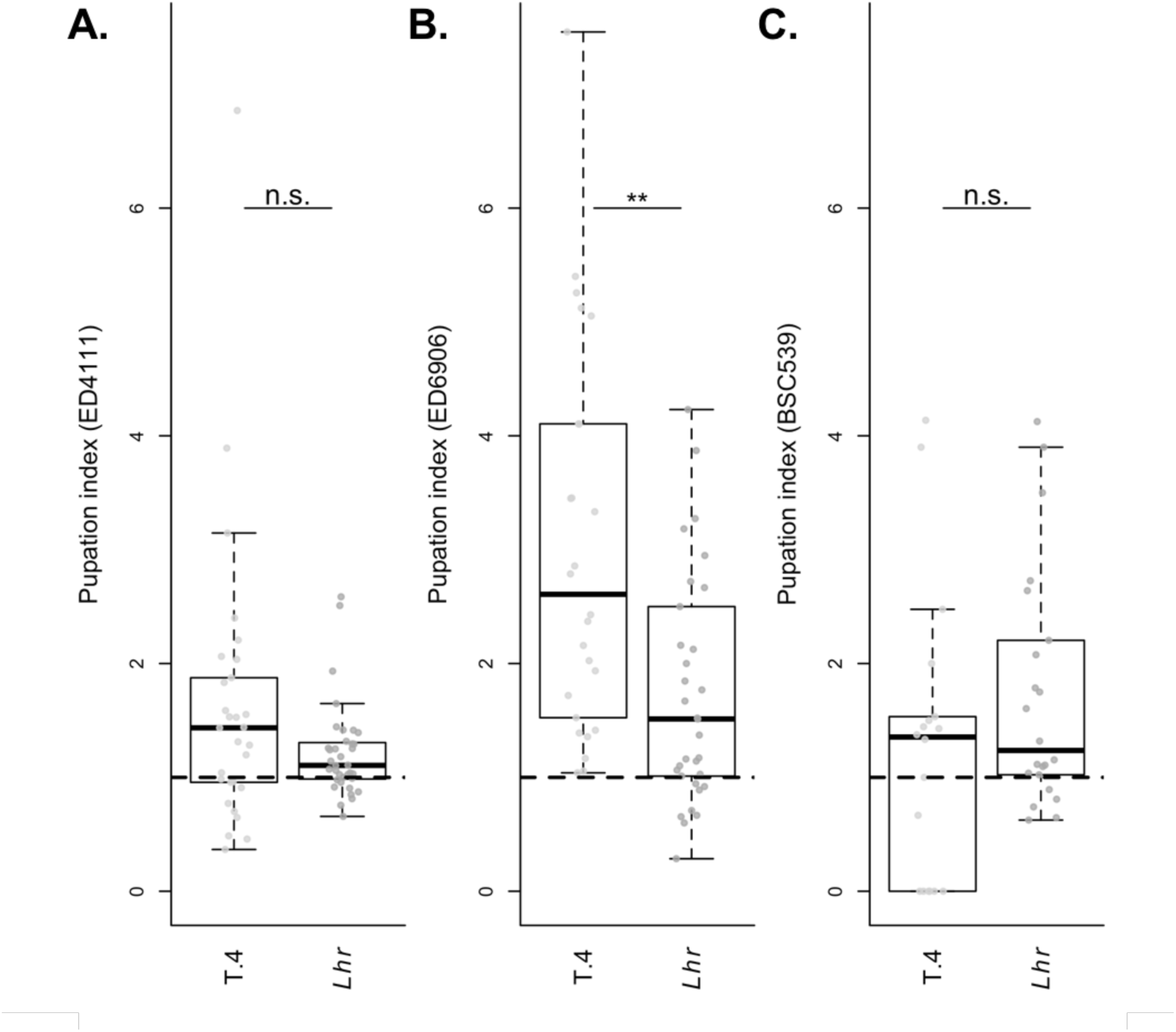
Pupation site choice behavior for additional significant hybrid deficiencies. Pupation indices for **A.** Df(1)ED411 (N=29-36), **B.** Df(1)ED6906 (N=26-33), and **C.** Df(1)BSC530 (N=18-22). Each of the three deficiencies displayed a median pupation index significantly greater than the average (0.88) when crossed to *D. simulans* (*Lhr*) after correcting for multiple comparisons. However, when each was crossed to *D. melanogaster* (T.4), the pupation index was still significantly elevated, indicating that the behavior of these flies is impacted by the deficient region in general, rather than the *Lhr* genotype it reveals.

**Fig S5.**
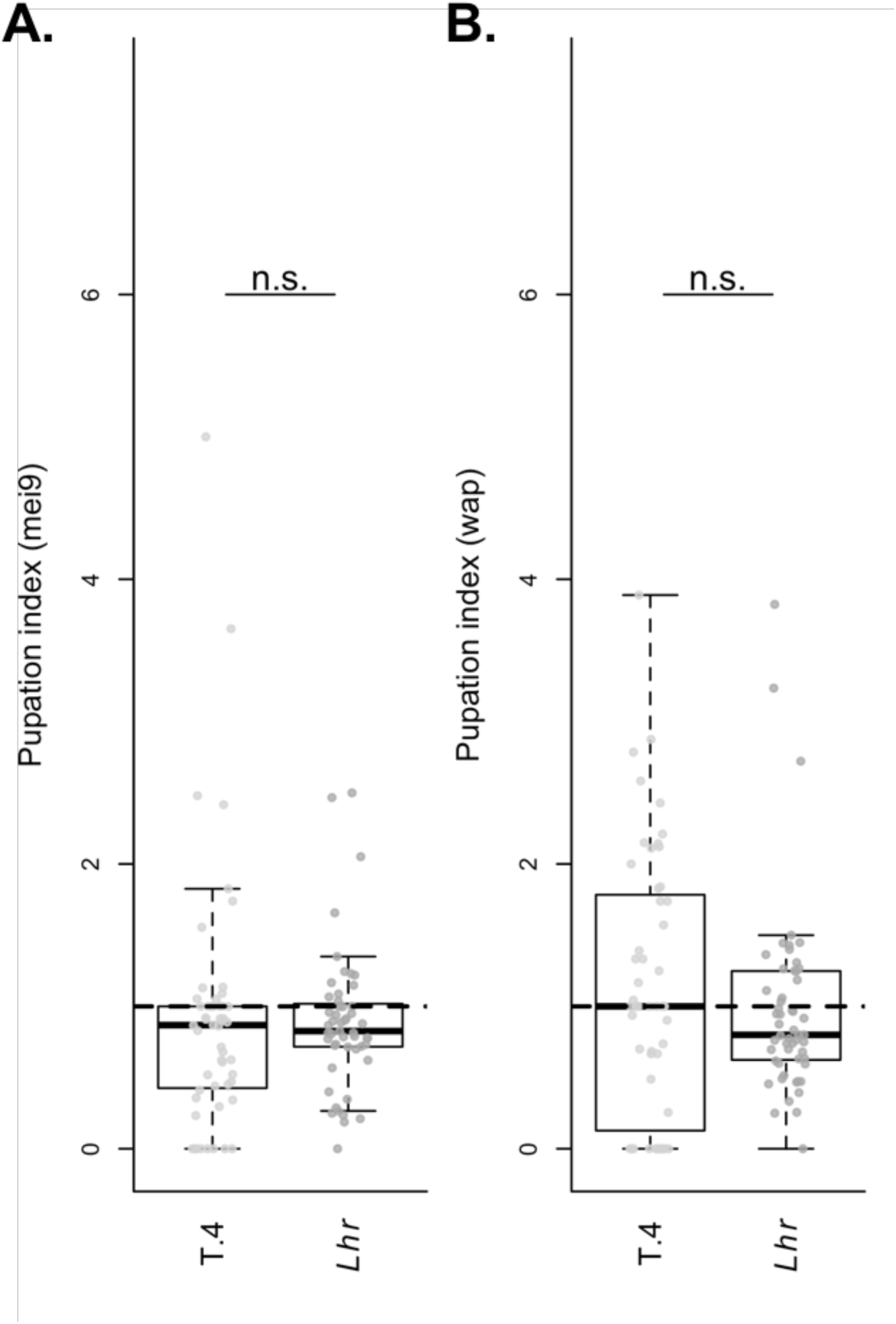
Pupation behavior for knockout strains crossed to both *D. melanogaster* and *D. simulans*. **A.** The pupation indices of a *mei9* gene disruption crossed to *D. melanogaster* (T.4, N = 52) and *D. simulans* (*Lhr*, N = 51). **B.** The pupation indices of a *wap* gene disruption crossed to *D. melanogaster* (T.4, N = 51) and *D. simulans* (*Lhr*, N = 55).

**Fig S6.**
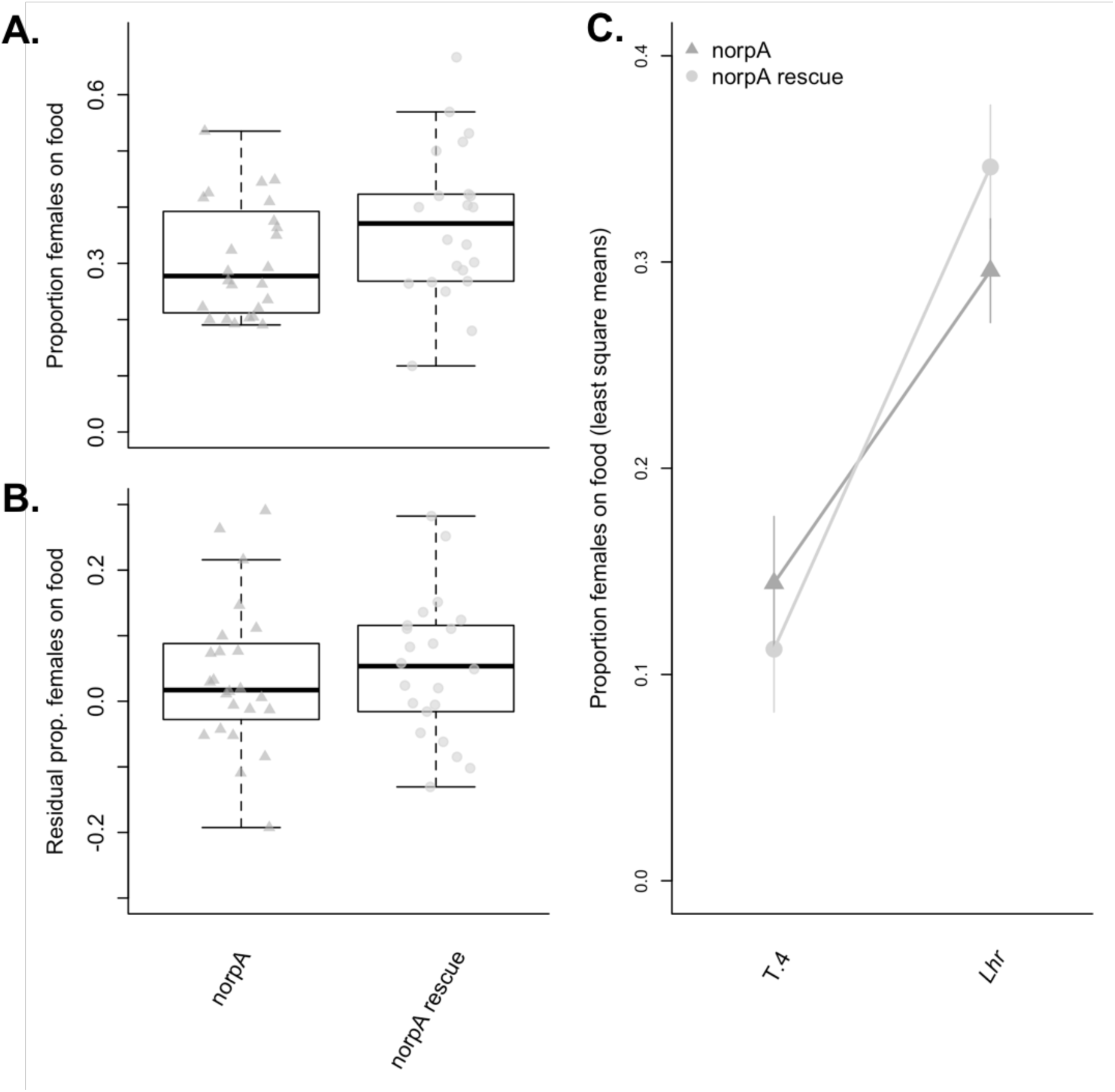
Pupation behavior for norpA knockouts and control. **A.** The proportion of females that pupated on the food is shown for the *norpA* mutant allele (norpA) and the *norpA* rescue allele (norpA rescue) when crossed to *D. simulans* (*Lhr*). There was no significant difference in pupation behavior between the knockout and the rescue allele (p=0.08). **B.** Pupation behavior for *norpA* knockout and *norpA* rescue allele female hybrids (crossed to *Lhr*) after controlling for density. Vials were adjusted for density using the residuals from a regression between the proportion of females in a vial that pupated on the surface of the food and the total number of individuals in the vial. There was no significant difference between the knockout and rescue allele after controlling for density (p=0.52). **C.** The proportion of females that pupated on the food when the *norpA* mutant allele (dark grey, triangle) and the *norpA* rescue allele (light grey, circle) were crossed to *D. melanogaster* (T.4) and *D. simulans* (*Lhr*). Shown are the least square means (± 1 SE) from an ANOVA with “species”, “*norpA* allele” and their interaction as fixed effects. The actual analysis was conducted using residuals to control for density, and the “*norpA* allele” x “species” interaction term was removed from the model as it was not significant (p=0.37).

**Fig S7.**
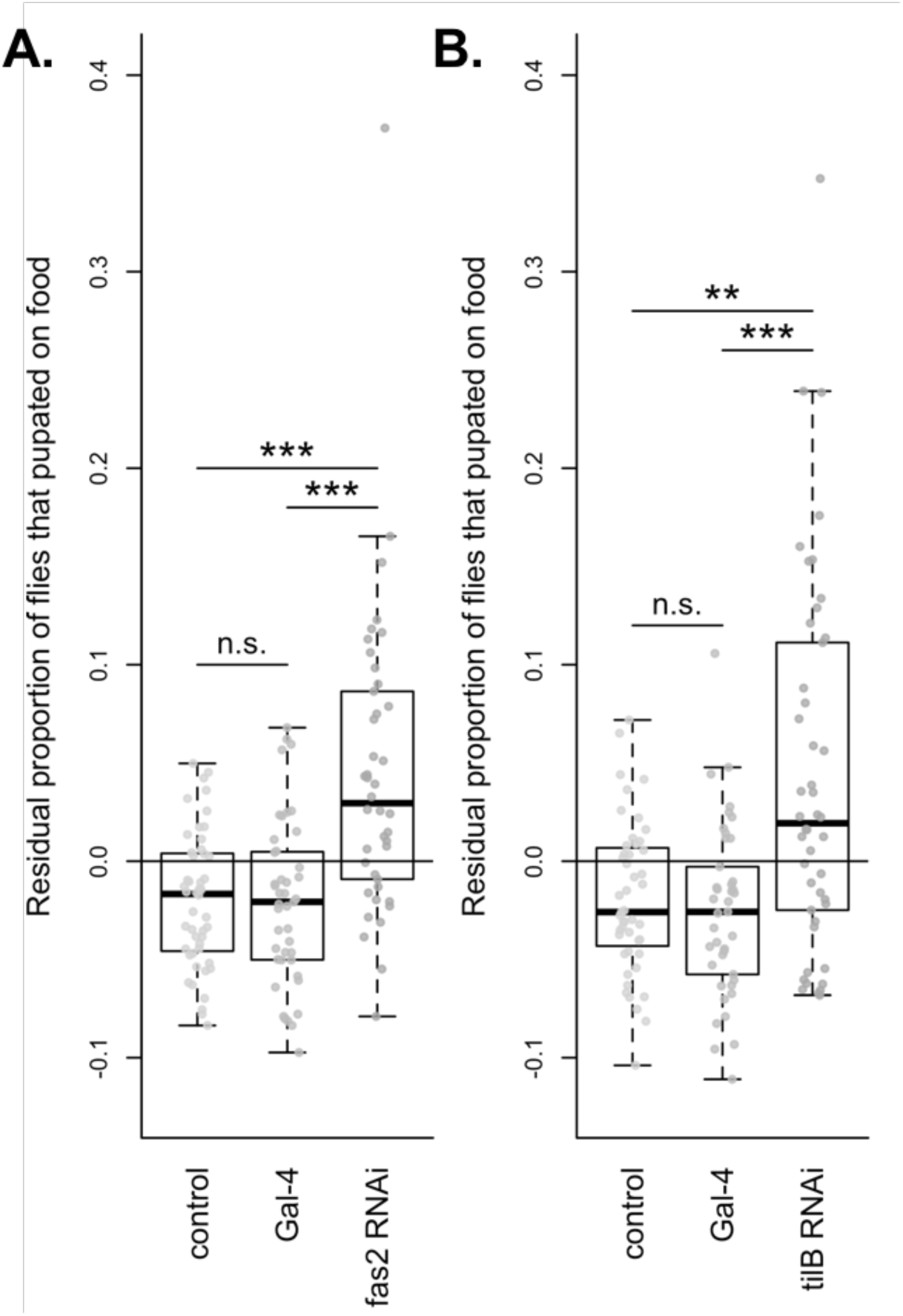
Pupation site choice behavior for *Fas2* and *tilB* RNAi crosses after controlling for density. **A.** The results of pan-neuronal knock down of the *Fas2* transcript via RNAi. Shown are the mean residuals from a regression between the proportion of individuals that pupated on the surface of the food and the total number of individuals in the vial for the control cross, the Gal-4 line, and RNA interference cross. Asterisks denote significance after correcting for multiple comparisons (* = p < 0.05, ** = p < 0.01, ** = p<0.001; N = 42-48). **B.** The results of pan-neuronal knockdown of the *tilB* transcript via RNAi. Shown are the mean residuals from a regression between the proportion of individuals that pupated on the surface of the food and the total number of individuals in the vial for the control cross, the Gal-4 line, and RNA interference cross. Asterisks denote significance after correcting for multiple comparisons (* = p < 0.05, ** = p < 0.01; N = 41-47).

**Fig S8.**
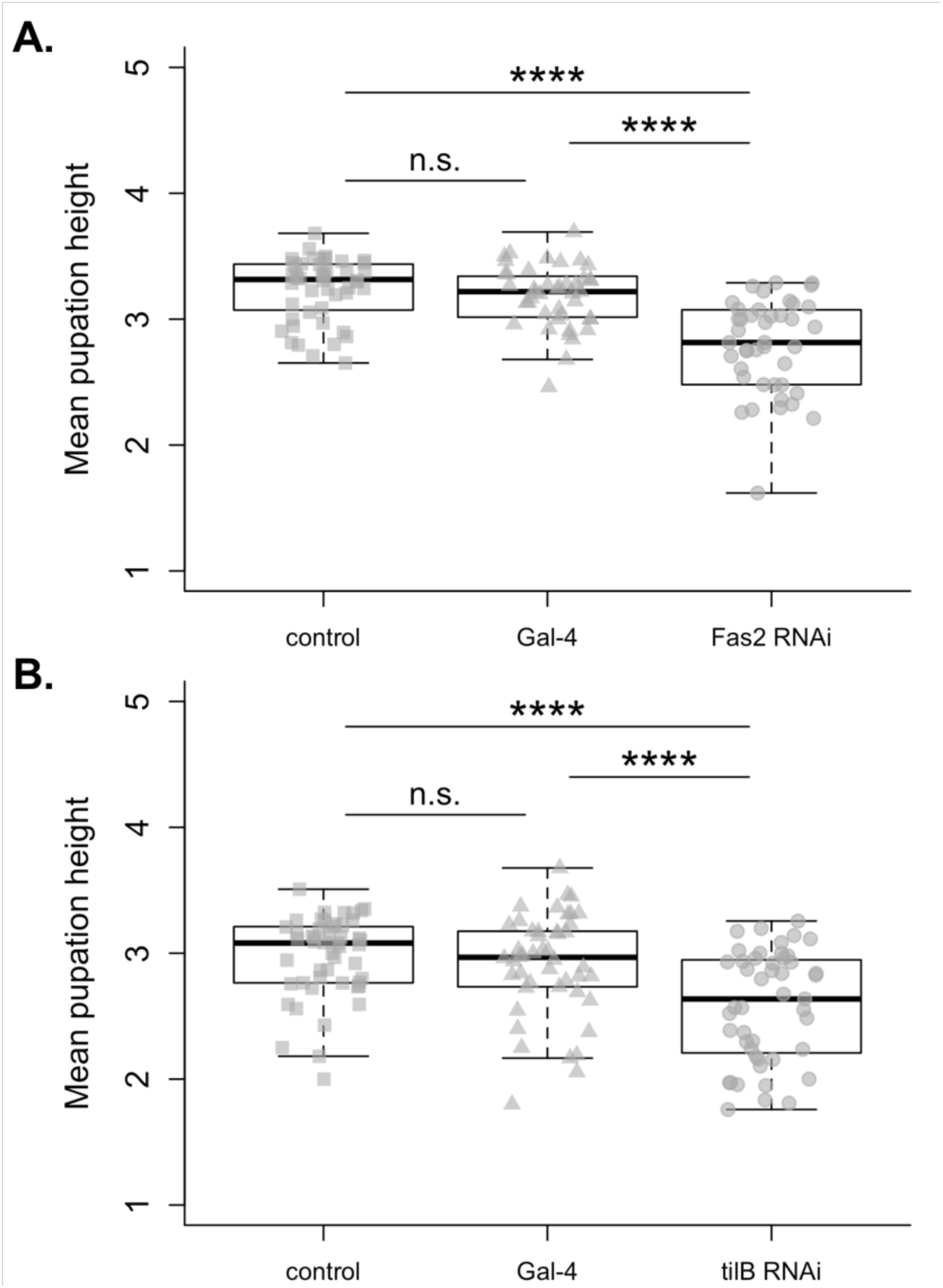
Pupation site choice behavior for *Fas2* and *tilB* RNAi crosses, measured as average pupation height within a vial. Higher pupation heights indicate individuals pupated further from the food. **A.** The results of pan-neuronal knockdown of the *Fas2* transcript via RNAi. The average pupation height of individuals found on the food are shown for the control cross, the Gal-4 line, and RNA interference cross. Asterisks denote significance after correcting for multiple comparisons (**** = p<0.0001; N = 42-48). **B.** The results of pan-neuronal knockdown of the *tilB* transcript via RNAi. The average pupation height of individuals found on the food are shown for the control cross, the Gal-4 line, and RNA interference cross. Asterisks denote significance after correcting for multiple comparisons (*** = p<0.0001; N = 46-45).

**Fig S9.**
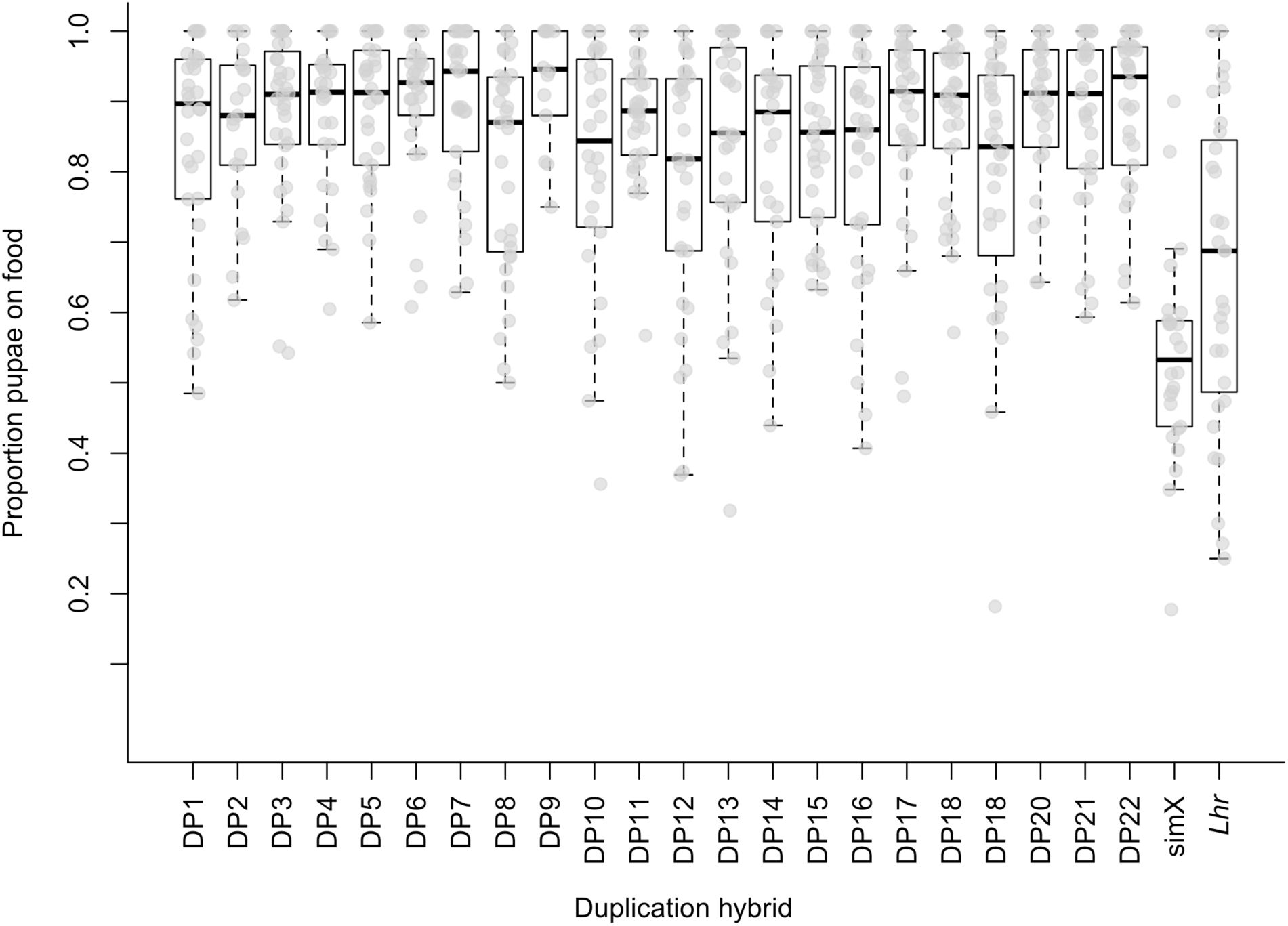
Duplication Screen. The results of a duplication hybrid screen. Each duplication hybrid was created by crossing a DP(1;Y) *D. melanogaster* female, which carries a unique segment of the X chromosome attached to the Y chromosome in an XXY background, to an *Lhr D. simulans* male. The resulting hybrid males have a complete *D. simulans* X chromosome in addition to 1 of 22 segments of the *D. melanogaster* X chromosome translocated to the Y chromosome. Females are largely inviable. In this way, we tested for dominant *D. melanogaster* X loci underlying the interspecific pupation difference. We scored pupation behavior following the same methods used for deficiency and knockout hybrids, and measured the proportion of pupae on the food for each duplication hybrid. Shown for comparison are the values for males from the *D. simulans* parent strain (*Lhr*) and hybrid males with a *D, simulans* X chromosome (simX). Interestingly, all of the duplication hybrids had significantly more males pupating on the food surface than either their *D. simulans* parent strain or typical simX hybrids that are not heterozygous for a region of the X. From this, we conclude that making males heterozygous over broad regions of the X chromosome results in behavioral or developmental inconsistencies that leave larvae unable to climb the vial walls to pupate. Information about these Y-linked X duplications can be found in Table S5.

